# Unsupervised Restoration of a Complex Learned Behavior After Large-Scale Neuronal Perturbation

**DOI:** 10.1101/2022.09.09.507372

**Authors:** Bo Wang, Zsofia Torok, Alison Duffy, David Bell, Shelyn Wongso, Tarciso Velho, Adrienne Fairhall, Carlos Lois

**Author notes:** These authors contributed equally to this work (Bo Wang, Zsofia Torok, Alison Duffy). Senior authors (Tarciso Velho, Adrienne Fairhall, Carlos Lois). (C.L.), (B.W).

## Abstract

Reliable execution of behaviors requires that brain circuits correct for variations in neuronal dynamics. Genetic perturbation of the majority of excitatory neurons in a brain region involved in song production in adult songbirds with stereotypical songs triggered severe degradation of their songs. The song fully recovered within two weeks, and substantial improvement occurred even when animals were prevented from singing during the recovery period, indicating that offline mechanisms enable recovery in an unsupervised manner. Song restoration was accompanied by increased excitatory synaptic inputs to unmanipulated neurons in the same region. A model inspired by the behavioral and electrophysiological findings suggests that a combination of unsupervised single-cell and population-level homeostatic plasticity rules can support the observed functional restoration after large-scale disruption of networks implementing sequential dynamics. In the model the sequence is restored through a parallel homeostatic process, rather than regrown serially, and predicts that sequences should recover in a saltatory fashion. Correspondingly, we observed such recovery in the songs of manipulated animals, with syllables that rapidly alternate between abnormal and normal durations from rendition to rendition until eventually they permanently settled into their original length. These observations indicate the existence of cellular and systems-level restorative mechanisms that ensure behavioral resilience.

---

Animal survival and reproduction requires reliable execution of behaviors. However, neuronal representations change over time, as a consequence of natural drift, or due to neuronal perturbations caused by trauma, disease, or aging^1,2^. What are the mechanisms that allow brains to maintain reliable behaviors over long periods of time or after neuronal loss? To investigate this question, we studied the zebra finch, a songbird that after learning a song as juveniles, produces stereotypical renditions of the song with minimal variability over several years. Songbirds have a series of brain nuclei dedicated for song learning and production called the song system^3^. HVC is the premotor nucleus in the song system that projects to the motor nucleus RA (robust nucleus of the arcopallium)^4^, involved in song production^3^. Previous experiments have shown that complete ablation of HVC abolishes song production^3^, but localized, partial lesions of HVC cause song degradation followed by recovery after different amounts of time^5–7^, suggesting that the song circuit is somewhat resilient. However, the precise behavioral, circuit and cellular level dynamics responsible for this resilience remain unknown. To investigate this question, we used genetic methods to selectively manipulate the activity of projection neurons in the HVC of adult finches and quantified the changes in the song as it degraded and recovered. In addition, we investigated the electrophysiological changes that occurred in this brain circuit after the genetic perturbation and created a data-inspired model to explore the cellular mechanisms underlying the observed behavioral and electrophysiological findings.

HVC projection neurons fire in an extremely sparse and precise manner when birds sing^8,9^. To explore how singing may be affected by disrupting the precise firing of these neurons, we expressed an ion channel to alter their electrical properties. Towards this goal, we bilaterally injected into both HVCs lentiviral vectors (LVs. **Fig. 1a**), which have been shown to selectively infect HVC projection neurons^10,11^ (**Extended Data Fig. 1**). To alter the electrical properties of HVC neurons, the LVs carried NaChBac, a bacterial voltage gated Na+ channel^12^. NaChBac expression in neurons perturbs their activity due to two features^13–15^: first, NaChBac is activated at membrane potentials in which the vertebrate Na+ channels are inactive; second, whereas the vertebrate Na+ channel depolarizations are several ms long, those of NaChBac expressing neurons last up to >1 s. The NaChBac transgene carried by the LVs contains the sequence of NaChBac fused in frame to GFP, which allows for visual identification of the infected neurons (**Extended Data Fig. 1**). Whole-cell patch clamp recording of HVC neurons in brain slices confirmed that GFP+ neurons displayed the characteristic NaChBac currents (**Fig. 1c and Extended Data 2a-c**).

**Figure 1.**
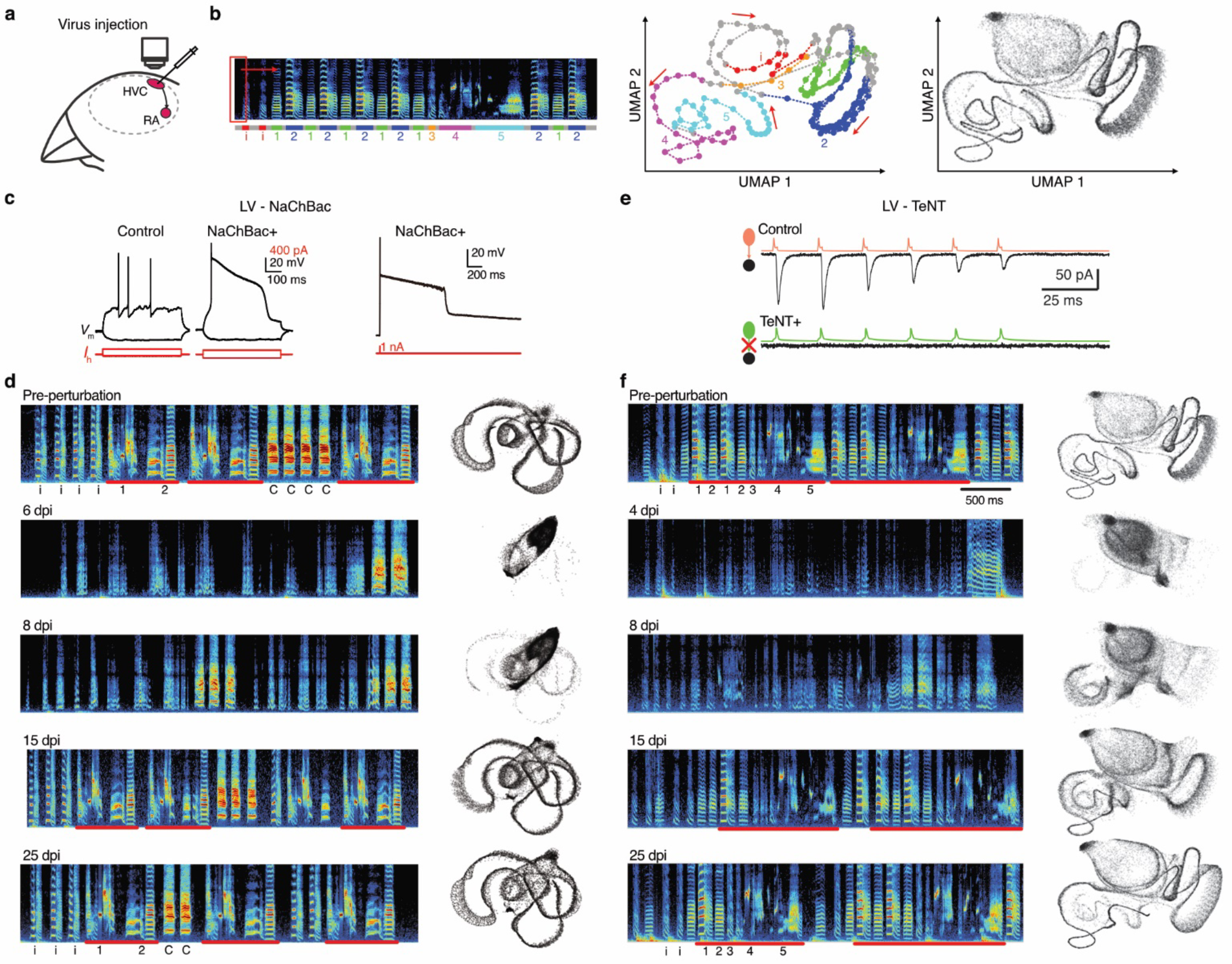
Song degradation and recovery after selective large-scale perturbation of excitatory neurons. **(a)** Schematic drawing illustrating the visual guided virus delivery into HVC (see Methods); **(b)** (Left) Spectrogram of a motif and the corresponding 2D projection (middle) using the UMAP algorithm from an unperturbed animal. The syllables are indicated by different colors and numbers in the spectrogram and UMAP plot. (Right) UMAP visualization containing ∼100 songs randomly sampled one day without perturbation; **(c)** Firing pattern of RA-projecting HVC (HVC(RA)) neurons with (NaChBac+) or without NaChBac (Control); **(d)** Example spectrograms and UMAP visualizations of the song of a bird injected with LV-NaChBac; **(e)** Example dual patch clamp traces demonstrating that expression of TeNT abolishes synaptic release from excitatory neurons in HVC; **(f)** Example spectrograms and corresponding UMAP visualizations of songs of a bird at different days post injection with LV-TeNT. The song motifs containing the syllables (12345) are marked by red lines and can be seen before perturbation and after recovery; ‘i’ stands for introductory note; ‘C’ stands for call.

For the first 2-3 days after viral injection there were no changes in the songs. Normal songs consist of repetitions of a so-called motif, which is made up of 3 to 7 syllables, depending on the individual. In unperturbed zebra finches, the acoustical structure of the syllables, the number of syllables per motif, and the duration of each motif are highly stereotypical, with minimal variations between renditions, even across months. In contrast, by 5 days after NaChBac injection the songs became highly irregular and bore no resemblance to their original songs.

To visualize the dynamics of the changes in these perturbed songs, we used Uniform Manifold Approximation and Projection (UMAP), a non-linear dimensionality reduction algorithm, to project the high-dimensional acoustic representation of hundreds of songs onto a 2D plane (see Methods and **Fig. 1b**)^16–18^. NaChBac delivery caused strong acoustic and temporal degradation across all syllables and greatly increased the variance between song renditions, which made it difficult to identify motifs (**Fig. 1d and Supplementary Audio 1-3**). However, at approximately 7 days post-injection (dpi) the song started to recover, and gradually regained structure. By 14 dpi, the songs of NaChBac animals were highly similar to their original songs and remained so for several months. To investigate how NaChBac expression caused song degradation, we performed whole-cell patch clamp recordings. As early as 48 hours after injection, NaChBac+ RA-projecting (HVC(RA)) neurons displayed the characteristic NaChBac current and abnormal long depolarizations. However, starting at 4 dpi, at the onset of song degradation, NaChBac+ HVC(RA) neurons had a substantial increase in inhibitory synaptic inputs leading to their silencing (**Extended Data Fig. 3**), as observed before in the mammalian brain^13,15,19^. These synaptic changes were also present at >25 dpi after the song had recovered. Thus, we hypothesize that the degradation of the song after NaChBac injection is primarily due to the eventual silencing of a large fraction of HVC projection neurons.

To investigate whether directly silencing HVC projection neurons would be sufficient to degrade the song, we delivered a LV carrying the light chain of tetanus toxin (TeNT). TeNT is an enzyme that cleaves synaptobrevin, a protein essential for the release of neurotransmitters. Thus, expression of TeNT in a neuron does not alter its electrical activity, but it abolishes its ability to communicate with its postsynaptic targets^20^. Paired whole-cell patch clamp recordings in brain slices showed that expression of TeNT in HVC(RA) projection neurons abolished their excitatory drive onto inhibitory neurons in HVC at all times, both during song degradation, and after recovery, indicating the TeNT permanently muted the projection neurons (**Fig. 1e and Extended Data Fig. 2d-f**).

**Figure 2.**
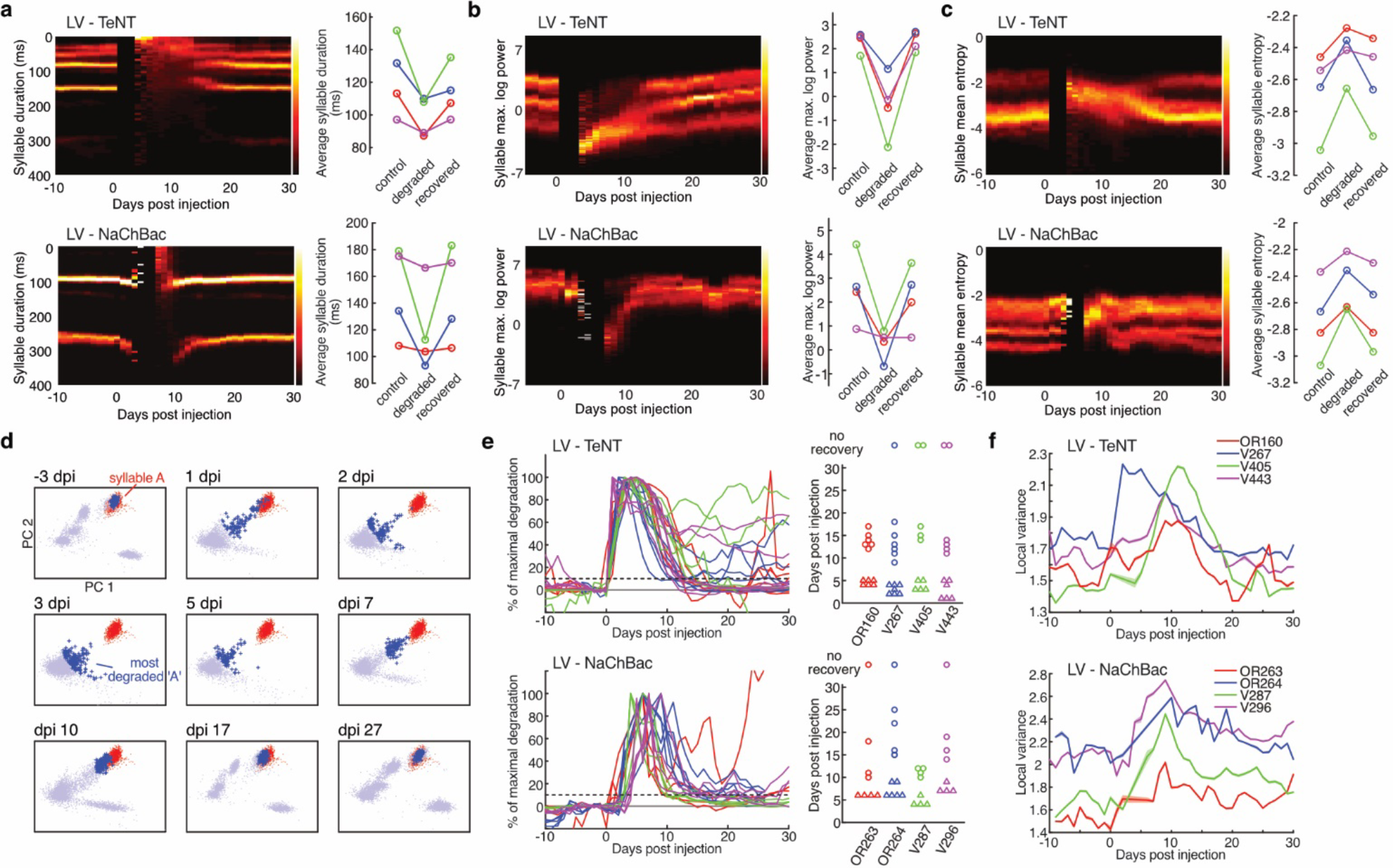
Dynamics of song degradation and recovery after large-scale perturbation of HVC excitatory neurons. **(a)** (Left) Density distribution of syllable durations day by day, for 2 birds either with LV-TeNT (upper) or LV-NaChBac (bottom). Note that the gap in the middle (between around 1-5 dpi) means the number of song bouts recorded that day was not enough for a meaningful analysis. (Right) Averaged duration of all syllables at three time points, before perturbation (control), when the song is most severely degraded (3-5 dpi), and when the song is fully recovered (25-30 dpi). Each line represents one animal; **(b)** (Left) Density distribution of the maximal log power of syllables day by day. (Right) Averaged syllable maximal log power at different time points. Animals with LV-TeNT on top and those with LV-NaChBac on bottom; **(c)** (Left) Density distribution of the mean entropy of syllables day by day. (Right) Averaged entropy of all syllables at different time points; **(d)** We tracked the change of song segments over time using acoustic features sampled from the spectrograms of the song segments (see also Methods). Here we show the trajectory of songs in the first 2 principal components of the acoustic space over a series of days after LV-TeNT injection. Each dot represents one original or degraded syllable. Red dots represent original renditions of syllable ‘A’ sung 1 day pre-perturbation. Blue dots represent song segments sung on each day. Dark blue dots represent the closest 2.5 % of song segments to the original syllable ‘A’ cluster on each day (measured using a k-nearest neighbor metric in the full acoustic parameter space), which we used to quantify the degradation of syllable ‘A’; **(e)** (Left) Plots of acoustic distance (see also Methods & Fig. S4) to each original syllable, normalized so that the peak of the curve represents 100% degradation. Curves in the same color were from the same bird. (Right) Triangles represent the day when each syllable reached peak degradation (TeNT 3.53 ± 0.39 vs NaChBac 6.25 ±0.66 dpi, p < 0.001, nested ANOVA), and round dots represent the day when each syllable achieved more than 90% recovery (13.71 ± 0.72 vs 14.80 ± 1.90 dpi, p > 0.05, nested ANOVA). Each column represents one bird; **(f)** Plots of the local variance across days.

Injection of LV-TeNT into HVC caused song deterioration and recovery with an overall similar pattern to that described for LV-NaChBac (**Fig. 1f and Supplementary Audio 4-6**), although songs started to degrade faster with TeNT (as early as 1 dpi). Lentiviral expression starts at ∼ 24 hours, TeNT is a powerful toxin, and a handful of TeNT molecules is sufficient to mute a neuron, thus this likely accounts for its rapid action. To investigate the dynamics of song changes caused by these genetic perturbations, we quantified multiple acoustic features of syllables of birds receiving either LV-TeNT or LV-NaChBac^21^. In both types of perturbation, degraded songs consist of shorter (**Fig. 2a**), weaker (**Fig. 2b**), and noisier (**Fig. 2c**) syllables than those of the original songs. Note, we use the term ‘syllable’ to refer to any continuous song segment in the degraded song, even if they are not stereotypical, and they bear little or no resemblance to the original syllables. We measured the acoustic distortion of songs by calculating the distance of disordered syllables to the original (pre-perturbation) syllables, normalized to the maxima, in a high dimensional acoustic space derived from the spectrograms (**Fig. 2d and Extended Data Fig. 4**. see also Methods) and plotted the trajectories of each syllable to show the timeline of dynamics of song degradation and recovery (**Fig. 2e**). We found that, both for LV-TeNT and LV-NaChBac animals, the acoustic distance to all original syllables increased drastically after the virus injection, but the onset and peak distortion occur earlier for LV-TeNT (3.53 ± 0.39 dpi) than for LV-NaChBac animals (6.25 ±0.66 dpi. **Fig. 2e**). Although the peak distortion happens later for NaChBac the time to recovery was comparable for the two types of manipulations (13.71 ± 0.72 vs 14.80 ± 1.90 dpi. **Fig. 2e**). The recovered song was very close to the original, but some localized aspects of the song changed permanently, including fine-scale acoustical structure, and loss of some syllables (**Fig. 1d**,**f**).

We also measured the rendition-to-rendition syllable variability across days (See Methods). For both NaChBac and TeNT, the peak of variability in syllable renditions occurred at around 10 dpi (**Fig. 2f**), approximately 4-6 days after the acoustic features of the songs were maximally degraded (**Fig. 2e**). By 20-25 dpi, the acoustic features were fully recovered, and the syllable variability returned to the pre-perturbation level (**Fig. 2f**). Importantly, the degradation of song was not due to toxicity or mechanical lesion, because no behavioral change was observed when the same amount of viral vector expressing either GFP or a dead-pore NaChBac was injected (**Extended Data Fig. 5**. N = 4). Finally, song recovery was not due to the restoration of normal firing activity in the NaChBac+ HVC(RA)neurons, because they still had the same amount of NaChBac current (**Extended Data Fig. 2a-c**) and highly increased inhibitory synaptic input at >25 dpi, 10 days after the song had recovered (**Extended Data Fig. 3**). Similarly, the muting of HVC(RA) neurons by LV-TeNT also persisted at >25 dpi, even after the song had recovered (**Extended Data Fig. 2a-c**). The similar behavioral outcomes after NaChBac and TeNT delivery suggests that the mechanisms by which these different genetic manipulations caused song degradation were due to the functional loss of a large percentage of HVC projection neurons, by direct muting caused by TeNT,or by homeostatic silencing in response to NaChBac.

**Figure 3.**
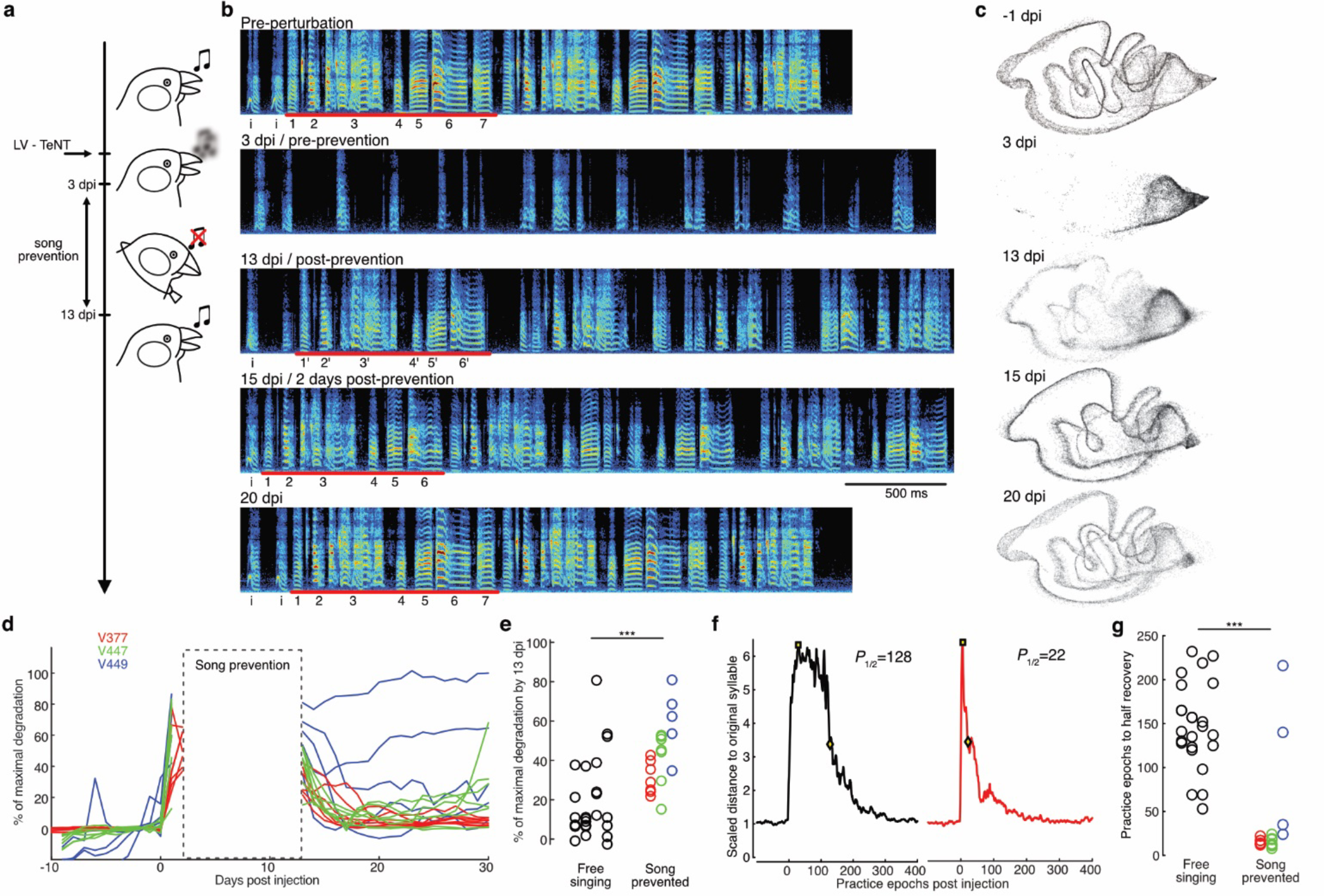
Partial recovery of degraded song without practice. **(a)** Schematic illustrating the experimental procedure of perturbation with LV-TeNT and song prevention; **(b)** Example spectrograms of a bird injected with LV-TeNT and prevented from signing for 10 days. On the first day after song prevention (“post-prevention”) the song was partially recovered, and syllables were good enough to match to those of unperturbed song (“pre-perturbation”); **(c)** UMAP visualizations of songs of the same bird shown in B; **(d)** Plots of distance to original syllables (normalized to maximum) of all birds prevented from singing (gray lines). Each color represents one bird. Prevented animals sang fewer than 8 songs per day versus more than 5,000 songs per animal per day for the freely singing animals during the 10-day prevention period (marked by a gray square); **(e)** At 13 dpi the song of freely singing birds had mostly recovered (20.4 ± 6.3%of the maximal degradation remaining. At this time, when song-prevented birds were allowed to sing they also showed significant recovery (44.2 ± 10.4% of the maximal degradation remaining) p < 0.001, Nested ANOVA; **(f)** Example plots of syllable recovery vs number of practice epochs with or without prevention; **(g)** Group data of the number of practice epochs to reach half recovery of each syllable, each column is one bird. Free-singing, 149.4 ± 12.4, N = 4; Song-prevented, 44.7 ± 29.5, N = 3. ***, p < 0.001, nested ANOVA.

Is the process of song recovery after HVC perturbation similar to the way young animals learn the song? Juvenile finches listen to the song of their fathers, memorize it, and attempt to copy it by continuous trial and error over 2-3 months and tens of thousands of practice renditions. The juvenile song is initially unstructured, until it eventually becomes a faithful copy of the father’s song, and it “crystallizes” into a stable motif with minimal variability^22^. To investigate if song recovery after HVC perturbations also required practice, we prevented LV-TeNT animals from singing during the two-week period when the recovery usually occurs, and after this time, they were allowed to sing freely (**Fig. 3a and Extended Data Fig. 6**). Surprisingly, the very first song renditions sung on the first day after prevention were already highly similar to the original pre-perturbation song (**Fig. 3b-g**). After singing a few dozens of renditions within the same day, the songs recovered to the same degree as in animals that had sung thousands of renditions for two weeks after viral injections. These observations indicate that much of the song restoration after perturbation could occur without practice and suggest that offline mechanisms could enable recovery in an unsupervised manner.

What are the cellular mechanisms by which the brain restores the precise execution of the song after such drastic perturbations? Both LV manipulations (NaChBac and TeNT) reduced the number of functioning HVC projection neurons. We hypothesized that the recovery may depend on changes among other neurons that were not genetically perturbed (“unmanipulated neurons”) within the same HVC. We performed whole-cell recordings to measure the synaptic and intrinsic properties of unmanipulated (GFP-negative) HVC(RA) neurons after LV-TeNT. These neurons showed no change in their intrinsic excitability or inhibitory synaptic inputs (**Extended Data Fig. 7**). However, they recruited a substantially higher level of excitatory synaptic inputs than neurons in unperturbed animals, as revealed by a much higher frequency of mEPSC (193.0 ± 31.5 %. **Fig. 4a-c**), suggesting that presynaptic mechanisms, such as formation of new synapses, activation of silent synapses, or strengthening of preexisting synapses might be responsible for these changes. Similar synaptic changes were observed in manipulated (TeNT+) neurons as well; however, because TeNT irreversibly muted these cells, it is unlikely that synaptic changes in the manipulated cells contributed to the recovery. To investigate whether these synaptic changes are caused by local perturbation of neuronal activity within the same region, we injected the virus unilaterally just in one HVC. We detected synaptic changes in unmanipulated neurons in the injected hemisphere, but not in the contralateral (unmanipulated) hemisphere (**Extended Data Fig. 8a**,**b**). This suggests that the observed synaptic changes are induced by local phenomena restricted to the HVC that was perturbed.

The excitatory inputs received by HVC inhibitory neurons in LV-TeNT animals were initially greatly reduced, consistent with a reduction of the synaptic output from the projection cells expressing TeNT. Dual whole-cell recordings between HVC(RA) projection neurons and interneurons indicated that expression of TeNT leads to permanent muting of the infected cells (**Extended Data Fig. 2**). However, these excitatory inputs to inhibitory neurons eventually recovered to a level that was indistinguishable from control animals (**Extended Data Fig. 8c**,**d**), suggesting that the recovery of the inhibitory tone was due to changes in unmanipulated neurons. Thus, our findings show that alongside song degradation and following recovery, the HVC network is drastically restructured.

We used modeling to explore how synaptic plasticity mechanisms could contribute to unsupervised recovery of network activity after perturbation and to account for the observed synaptic changes. Song recovery in animals that were prevented from singing suggests that plasticity mechanisms are initially largely driven locally within HVC rather than by behavioral feedback from practice. We sought to determine which mechanisms are consistent with restoration of the sequence as well as recapitulation of our physiological findings on synaptic strength changes. We modeled HVC as an excitatory-inhibitory (E-I) network, with HVC(RA) neurons connected to each other in a feedforward, polysynchronous chain and recurrently connected to HVC(INT) neurons (**Fig. 4d**)^9,23,24^. We then inactivated varying fractions of the HVC(RA) population (**Fig. 4d**) to mimic the silencing of neurons caused by TeNT expression. To explore mechanisms that may enable recovery, we first implemented spike-timing dependent plasticity (STDP) between excitatory neurons (E→E), which is integral to many models of sequence self-organization^23,25–27^. In addition, implementing STDP between excitatory and inhibitory neurons (E→I) enabled the rebound of input to inhibitory neurons observed in our experiments. However, we found that STDP alone did not reliably restore sequential activity (**Extended Data Fig. 9**), as the perturbed chain lacks activity in both pre- and postsynaptic neurons required to strengthen synapses, and further did not produce the overshoot in excitatory connectivity into HVC(RA) neurons observed by our electrophysiological measurements.

As STDP alone was insufficient to restore the sequential dynamics in our model of HVC, we next considered whether cell-autonomous homeostatic mechanisms based on changes on either intrinsic excitability or synaptic inputs may enable recovery of the network. Our recordings revealed that unmanipulated HVC(RA) neurons did not change their intrinsic excitability post-perturbation (**Extended Data Fig. 7**), but they displayed changes in their synaptic inputs. Thus, we added a cell-autonomous homeostatic rule into our model, based on scaling the excitatory synaptic inputs to individual excitatory neurons to maintain their firing rates (**Fig. 4e**). The implementation of this rule is consistent with reports that found activity-dependent synaptic homeostasis in other circuits with sequential dynamics^28^. We found that adding this form of cell-autonomous synaptic homeostasis in excitatory neurons reliably restored sequential network activity (**Fig. 4f,g**). However, in models employing STDP alone or both STDP and cell-autonomous synaptic homeostasis, the recovered HVC outputs were reduced in proportion to the percentage of neurons inactivated, thereby drastically weakening the drive to downstream regions (**Fig. 4h**). Additionally, these models did not reproduce the overshoot in excitatory synaptic input to unmanipulated excitatory neurons as revealed by our experiments (**Fig. 4c,g**).

One potential mechanism by which HVC might restore its output to downstream targets is the recruitment of neurons that initially do not participate in the sequential dynamics, defined here as “silent” neurons (**Fig. 4i**). Multiple experiments have indicated that a large fraction of HVC(RA) neurons do not fire during song, suggesting a possible redundant role^8,29,30^. We hypothesized that the presence of such silent neurons--assumed to be HVC(RA) neurons connected within the network, but whose inputs are subthreshold--might provide additional resilience by allowing the sequential dynamics to be partially carried by newly recruited HVC neurons when active constituents of the network fail. While such shifts in representation could be due to the loss of inhibition onto silent cells following the loss of excitatory neurons, this picture of recovery is inconsistent with our experiments in that it does not require the large increase in the excitatory inputs onto HVC(RA) neurons that we observed in our recordings. We therefore hypothesized that silent neurons may be recruited into the sequence through a form of population homeostasis, for which there is an emerging body of support^31–34^. In our model, we implemented one potential mechanism by which such population homeostasis can be achieved: synaptic scaling based on the activity-dependent release of secreted factors (**Fig. 4j**), such as BDNF and TNFα, which have been shown to regulate local network activity in a non-cell-autonomous manner^35–38^. While local population homeostasis alone enabled reliable recovery of sequences (**Extended Data Fig. 10**), the recruitment of previously silent neurons enabled the most complete recovery consistent with our experiments in that the total output of the network was restored and E→E synaptic inputs increased by ∼100% (**Fig. 4k-m**). Furthermore, the dynamics of the sequence in terms of numbers of participating neurons and their temporal resolution most closely resembles the pre-perturbation state. Finally, the recruitment of silent neurons (either mature cells already present or potentially new neurons produced during adult neurogenesis^7^) may initially add noise to song production (**Fig. 4k**), consistent with the observation of an increase in the song’s variability, peaking at ∼10 dpi, approximately 4-6 days after the acoustic features of the songs were maximally degraded (**Fig. 2f**).

**Figure 4.**
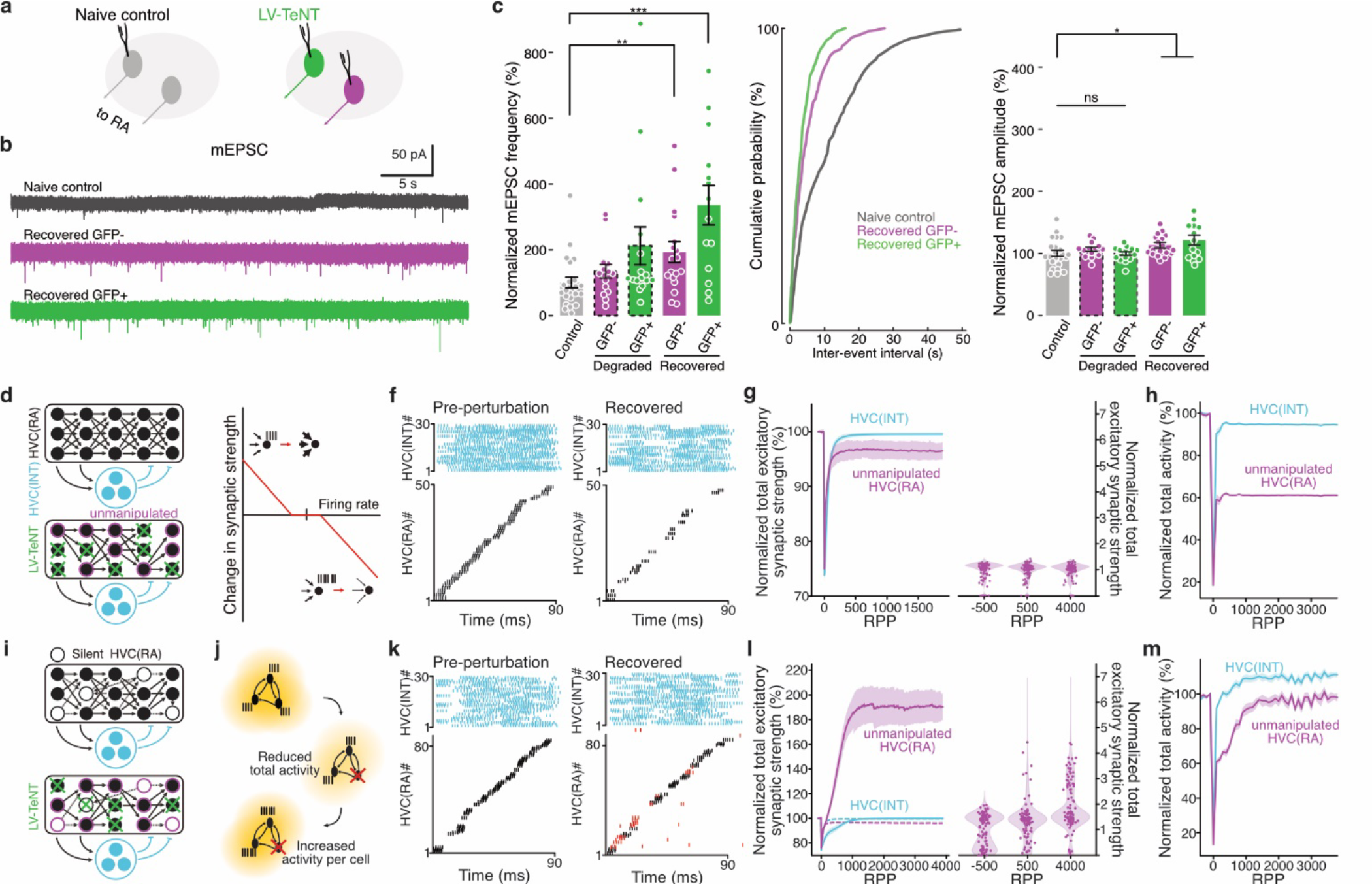
Population-level homeostatic plasticity and recruitment of silent neurons in a network model contribute to the recovery of sequential activity. **(a)** Schematic illustrating whole-cell recordings from HVC(RA) neurons in birds injected with LV-TeNT or naive controls; **(b)** Example traces of mEPSC recordings; **(c)** Group data of mEPSCs. “Degraded” indicates recordings at 5 dpi when the song was degraded. “Recovered” indicates recordings at 25 dpi, after the song had fully recovered. mEPSC frequency: Control, 9.7 ± 1.6 min^-1^, N = 23/4; Degraded GFP-, 13.1 ± 0.2 min^-1^, N = 15/3; Degraded GFP+, 20.6 ± 0.3 min^-1^, N = 16/4; Recovered GFP-, 18.8 ± 0.3 min^-1^, N = 18/4; Recovered GFP+, 32.7 ± 0.5 min^-1^, N = 14/4. mEPSC amplitude: Control, 17.2 ± 0.9 pA; Degraded GFP-, 18.3 ± 0.6 pA; Degraded GFP+, 17.1 ± 0.5 pA; Recovered GFP-, 19.5 ± 0.7 pA; Recovered GFP+, 20.8 ± 1.4 pA. *, p < 0.05. **, p < 0.01. ***, p < 0.001, One-way ANOVA & student’s t-test. Error bars represent s.e.m.; **(d)** Schematic diagram of the neuronal organization in the model; **(e)** Schematic illustrating the single-cell homeostatic plasticity rule implemented; **(f)** Spike raster plots showing the sequential dynamics generated by HVC neurons before perturbation and after recovery (with only single-cell homeostatic plasticity implemented); **(g)** (Left) Plot of the normalized total excitatory synaptic input per neuron against renditions. RPP, rendition(s) post perturbation. (Right) Scatter plot of the normalized total excitatory synaptic input received by each HVC(RA) neuron at three time points; **(h)** Plots of the normalized total firing activity of all functioning neurons; **(i)** Schematic diagram illustrating that initially inactive neurons were recruited into the network; **(j)** Schematic diagram illustrating that a population-level homeostatic plasticity, governing the summed firing activity of all neurons, was also implemented in the model; **(k)** Spike raster plots showing the sequential dynamics generated by HVC cells before perturbation and after recovery; **(l)** (Left) Plot of the normalized total excitatory synaptic input per neuron against number of renditions. Dashed lines are adapted from panel g for comparison. (Right) Scatter plot of the normalized total excitatory synaptic input received by each HVC(RA) neuron at three time points; **(m)** Plots of the normalized total firing activity of all functioning neurons. Shadows in g and i represent standard deviation, and in h and m standard error is shown.

We use modeling, beginning from standard assumed architectures for HVC, to explore the potential roles of different plasticity rules and to seek conditions that lead to agreement with experimental data. The core of our model is that the sequence is more reliably restored through a parallel homeostatic process, in contrast to serial regrowth through a timing-dependent process (**Fig. 5a and Extended Data Fig. 9**). A prediction of the model is that the sequence should recover in an abrupt fashion: all links are repaired simultaneously, so that by the time any broken link is restored, all other links tend to have recovered (**Fig. 5b-d**). Correspondingly, we found many examples of such saltatory recovery in the song, whereby syllables show a bimodal temporal distribution (**Fig. 5e-h**), with syllables rapidly alternating between two durations from rendition to rendition until eventually they permanently recover their original duration. Syllables with these bimodal temporal distributions are only present in renditions that include song segments that bear acoustic resemblance to the original syllables. This suggests the existence of a remaining chain “skeleton” that supports production of the correct syllable on occasional renditions until the sequence’s strength fully recovers. A further implication of these observations and the recovery set out by the model is that the skeleton can act as a scaffold upon which the homeostatic plasticity can rebuild the dynamics in an unsupervised manner. As this redundancy can exist very ‘locally’ in the chain (i.e. only between adjacent links) it is still consistent with an extremely sparse network, and may be a key contributor to resilience in HVC.

**Figure 5.**
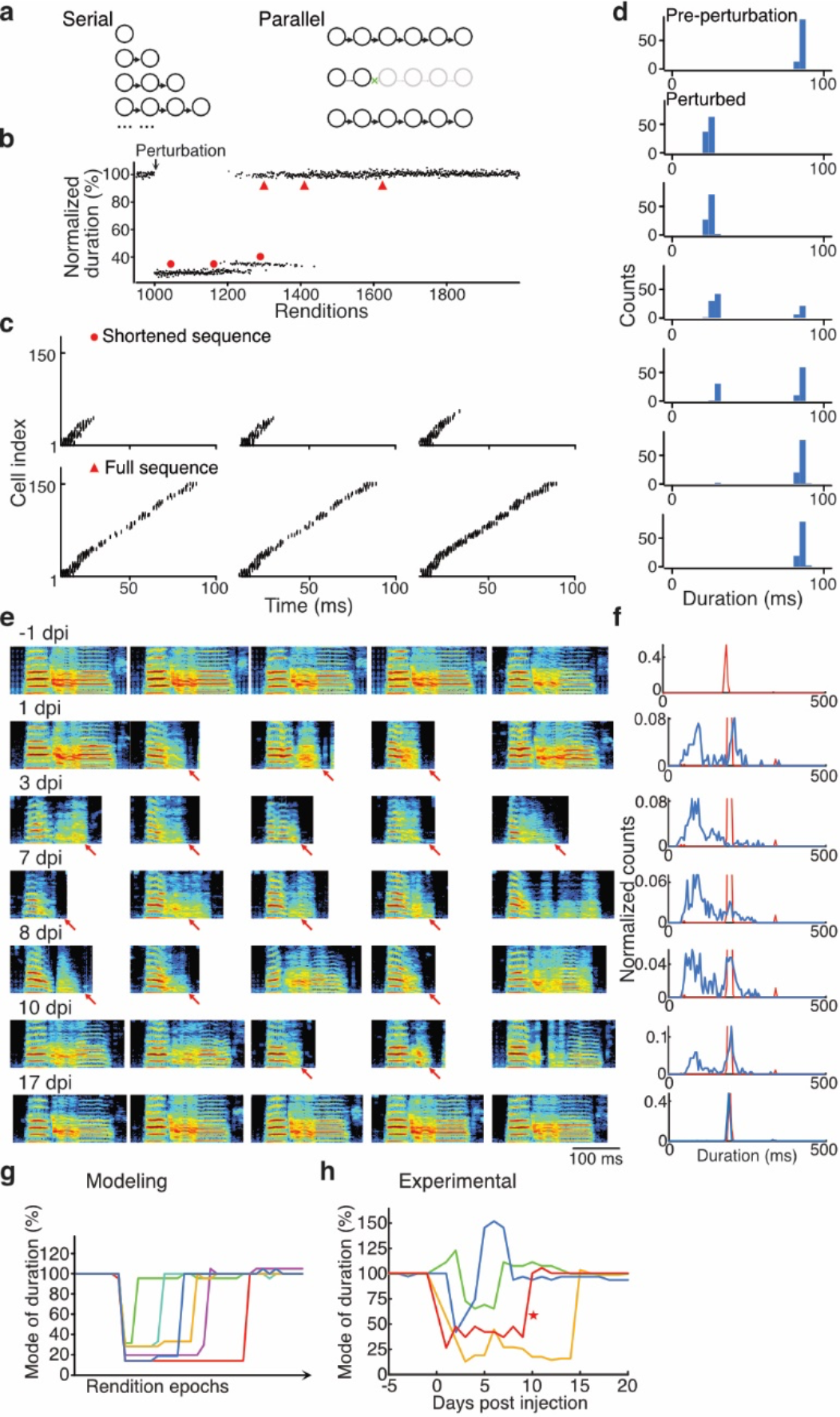
Saltatory recovery of syllable duration. **(a)** Schematic diagrams showing two types of potential circuit recovery mechanisms. (Left) Serial - the recovery of sequential firing requires building of feedforward synaptic chains step by step from the breaking point, such that the duration should regrow continuously. (Right) Parallel - all links in a sequence are repaired simultaneously, so that the recovery of the full sequence can be abrupt, when the broken links are fixed; **(b)** Plot of the normalized duration of modeled sequential firing against number of renditions. Note that the recovery of the full syllable duration was not continuous, but instead it occurred by a sudden leap from a shortened state; **(c)** Example raster plots showing the sequential spiking of modeled HVC(RA) neurons, picked from multiple time points indicated by red dots (shortened sequence) or triangles (full sequence) in panel b; **(d)** Probability density distributions of the durations of the modeled syllables at different times (before perturbation, during the degraded period, during the recovery period, and after full recovery), ordered chronologically. Note the bimodal distribution of the duration of a single syllable during the recovery phase; **(e)** Example spectrograms picked from the nearest k neighbors of one syllable at multiple days before and post injection of LV-TeNT. Red arrows mark the shortened/truncated syllables found between 1 to 10 dpi; **(f)** Probability density distributions of the duration of the k neighboring syllables, ordered so that each row of panels in e and f is from the same day. Note the bimodal distribution of durations for a single syllable found during the period of song degradation and recovery; **(g)** Plots of the mode of the duration of the modeled sequences against rendition epochs, showing that the recovery of sequence duration occurs in a saltatory, rather than in a continuous, manner; **(h)** Plots of the mode of duration of actual song syllables, which shows a saltatory recovery similar to that predicted by the model. The red curve marked with a star is made from the syllable shown in panels e and f.

The precise recovery of a complex learned behavior from extensive perturbations to a brain region essential for its production indicates the presence of cellular and systems-level restorative mechanisms that operate locally and in an “offline”, partially unsupervised manner. Song restoration occurred even when animals were prevented from singing during the recovery period, which indicates that the restoration mechanisms were mostly guided by neuronal activity without any information regarding how these cellular changes may improve behavior. Song recovery was accompanied by increased excitatory synaptic inputs to unmanipulated neurons within the same brain region. Based on these findings we modeled sequential dynamics to test how plasticity mechanisms can support this restoration. The inclusion of population-level homeostasis in the model confers a unique form of resilience as it enables the recruitment of HVC(RA) neurons that did not initially participate in song dynamics, providing a potential role for these inactive cells; neither STDP nor single-cell homeostasis can recruit these silent neurons into the active dynamics. The model predicts that the set of HVC(RA) projection neurons active in song would shift substantially following perturbation. Such population-level activity regulation could support naturally occurring “representational drift”, which has been observed in several brain regions wherein the qualitative nature of the sequential dynamics is preserved but the neurons that participate in the dynamics change^39–43^, potentially increasing circuit robustness^44,45^. Population-level homeostasis accounts for the experimentally observed increase in excitatory synaptic inputs to unmanipulated excitatory neurons as the song recovers. The observed upregulation of synaptic strength may have occurred through a random global process that could be later refined through activity-dependent plasticity. Instead, we find that the excitatory weight changes remain after the song has fully recovered in the experiment, suggesting, consistent with our modeling, that these changes represent a persistent network reorganization critical to the recovery process. Finally, the model recapitulated the experimental finding that restoration of sequential firing can happen offline (without feedback from practice) to result in song recovery, pointing to a largely unsupervised, circuit-level reorganization. We hypothesize that the potential for self-organized restoration of sequential dynamics may be key to enable such circuit mechanisms to support resilient behavior.

## Author contributions

All authors contributed to the conceptualization, methodology, and writing of the paper.

Investigation - Experiments: BW, ZT, SW, TV, CL

Investigation - Computational: AD, DB

Visualization: BW, ZT, AD, DB

Funding acquisition & Supervision: AF, CL

## Data and materials availability

All derived data in this study are included in this article. Raw datasets and computer codes are available from the corresponding authors on reasonable request.

## METHODS

### Animals

All procedures involving zebra finches are approved by the Institutional Animal Care and Use Committee of the California Institute of Technology. All birds used in the current study were bred in our own colony and housed with multiple conspecific cage mates of mixed sexes and ages until use for experiments. Before any experiments, adult male birds (>120 days post hatch (dph)) were single housed in sound isolation cages with a 14/10 hr light/dark cycle for >5 days until they were habituated to the new environment and started singing. Thereafter, birds were kept in isolation until the end of the experiment.

### Viral vectors

Lentiviral vectors were cloned using standard procedures and were produced and titrated as described previously^46^. All LVs contained the internal Rous sarcoma virus (RSV) promoter driving expression of different transgenes. LV-TeNT encoded the light chain of tetanus toxin fused to EGFP with a PEST domain in its C-terminus. LV-NaChBac encoded the open reading frame of NaChBac fused to EGFP.

### Stereotaxic injection

Birds were anesthetized with isoflurane (0.5% for initial induction, 0.2% for maintenance) and head-fixed on a stereotaxic apparatus. To inject a retrograde tracer in area X or RA, craniotomies were made bilaterally and fluorescent tracers (cholera toxin b 555 0.2%, fluoro-ruby 10%, or red RetroBeads, 100-300 nL) were injected through a glass capillary (tip size ∼25 μm) into the corresponding nuclei (coordinates from dorsal sinus in mm - area X: Anteroposterior (AP) 3.3-4.2, Mediolateral (ML) 1.5-1.6, Deep (D): 3.5-3.8; RA: AP 1.5, ML 2.4, D 1.8-2.1). To deliver virus into HVC, a second surgery was performed 7-10 days after retrograde tracer injection, by then HVC was strongly labeled by fluorescence and visible through a fluorescent stereoscope. Because LVs only diffuse a short distance from the injection site (∼100-200 µm), they were injected into multiple locations (up to 16 sites per hemisphere, ∼100 nL each) to deliver the transgenes into as many cells as possible throughout HVC. All injections in HVC were performed ∼20 nL/min to minimize physical damage. At the end of every surgery, craniotomies were covered with Kwik-Sil and the skin incision was closed with Gluture.

### Song analysis

Song analysis was performed using Matlab (Mathworks).

#### Song feature parameterization

Continuous audio recordings (44.1 kHz) were segmented into individual bouts manually and filtered to remove low frequency noise (cutoff frequency = 500 Hz). We used the open source Matlab software package, Sound Analysis Pro 2011 ^21^ to generate spectrograms and derive non-linear, time-varying song parameterizations. The time-varying features were: pitch, goodness of pitch, Wiener entropy, frequency modulation (FM), amplitude modulation (AM), amplitude, aperiodicity, mean frequency, and the maximum power in each quadrant of the frequency range 0-11 kHz (labeled power 1, power 2, power 3 and power 4). These features were computed across the entire bout every 1 ms. These time-varying features were the base from which various portions of the song were further characterized.

#### Syllable parameterization

A high-dimensional, acoustic parameterization of individual syllables was generated by then sampling a moving average (over 10 ms) of song features (pitch, goodness of pitch, mean frequency and entropy) at 10 points across the first 50 ms of each syllable. This resulted in a 40-dimensional song feature parameterization of each syllable. If a syllable was shorter than 50 ms, the feature vectors were padded at the end with zeros. This method allowed syllables to be compared in the same parametric acoustic space despite differences in duration.

#### Syllable segmentation

We identified syllables and silences within each bout by imposing thresholds on the time-varying, total log-power of the spectrogram. To consistently assign inter-syllable breath sounds as silence and to address minor variations in background noise during recordings across days, thresholds were defined by sampling from the background noise within each bout. We first applied a song threshold below which the recording window was defined as silence. This threshold was chosen to be high so that it would capture time windows between syllables in which soft breath sounds occur. The mean (µ) and standard deviation (σ) of the silent regions for that bout were computed from the middle portions of the silent windows. A new threshold was defined as: *T* = µ + *a* ∗ σ within that bout. The multiplier, *a*, was selected upon inspection of the stereotyped song structure for each bird and then applied across all recorded bouts across all days. We then performed a smoothing step wherein periods of silence less than 15 ms were converted to song segments. Song segments less than 20 ms were excluded from analysis. Segmentation was further checked by eye by random sampling across both stereotyped motifs and degraded songs. We then applied these parameters to the entire course of song recordings for each bird.

A note on terminology: we refer to song segments to indicate continuous periods of singing. In the unperturbed song, these song segments are termed syllables. Because this is a continuous recovery process, these terms sometimes overlap in our usage.

#### Syllable feature distributions

Syllable features (duration, log power, and entropy) were extracted from the syllable segmentation and parameterization processes. Distributions of syllable features were computed by normalizing each distribution of all syllable features within individual days such that the sum of each daily distribution over all binned durations equaled one. Distributions for individual days were then assembled into a matrix wherein the columns represented normalized distributions for individual days. This matrix was plotted as a heat map (Fig. 2a,b,c and Extended Data Fig. 5,e).

#### Acoustic distance trajectories

To quantify fluctuations in song, we computed k-nearest neighbor statistics of the acoustic parameterization space of song^47^. We computed the average k-neighbor distances of individual syllables from original, stereotyped syllables in the undistorted song. This distance was calculated from the first 20 principal components of the 40-dimensional, acoustic space described in Syllable feature parameterization. We defined a normative set of syllables from 1-3 days of recording pre-viral injection when the bird was singing undisrupted, stereotyped songs. This syllable set was labeled according to syllable identity. Syllable identity was defined using the Matlab clustering algorithm, dbscan. The dbscan clustering was performed on a reduced 2-dimensional acoustic space generated using the dimensionality reduction algorithm, Uniform Manifold Approximation and Projection (UMAP)^16,48^. The syllable assignments were cross checked by visual examination of a randomly selected subset of bouts and found to be in strong alignment with hand-marked syllable assignments. Syllables that did not cluster into distinct groups were excluded from this analysis. Each individual syllable was then assigned a k-nearest neighbor distance (k = 25) from the syllables in each normative syllable cluster, which we calculated as the average distance to the 25 closest syllables within the defined cluster.

As a measure of song degradation, we tracked the acoustic distance of the syllables which most closely resemble our original syllable set, defined within a local window of singing. We quantified the k-neighbor distance of the closest syllables to each original syllable cluster in a local window of consecutively sung syllables (N = 400 syllables; the 2.5% quantile of local k-neighbor distances to each syllable cluster) (Fig 2e,f; 3df; S4). When the song is highly distorted, syllables do not cluster into clearly defined groups, nor do they resemble original syllable types. This method allows us to track syllable distances even when syllables are highly distorted. We normalized the acoustic distance trajectories such that the peak acoustic distance was 100% distortion in order to compare the time course of recovery across birds and syllables.

#### Speed of recovery

We used the acoustic distance trajectories to measure the speed with which the song recovers. We measured the point of half recovery by tracking how much time and practice respectively are required post-perturbation to achieve a 50% recovery to the pre-perturbation baseline. 50% recovery is calculated relative to the point of maximum syllable distortion for each syllable individually. The acoustic trajectory of recovery is described in the above section. This measurement is shown for the speed of recovery as a function of practice in Fig 3f,g.

#### Song variability

As a measure of song variability, we tracked the local variability of syllables to other syllables that have been sung within a consecutive 400 syllable window (Fig. 2f). We quantified local variability as the average k-neighbor distance of each syllable to the closest 5 syllables within the local 400 syllable window. This measure of variability quantifies how different renditions are from one another, not how similar they are to the original song.

#### Continuous representation of bout trajectory

We generated continuous visualizations of bouts across the entire perturbation trajectory as shown in Figs. 1b,d,f; 3c, and Extended Data Fig. 5 ^17,18^. We randomly sampled 100 bouts from each day of recording to build a representative sample of the song over the course of the experiment. For each bout, we slid a 150 ms window in 3 ms steps along the bout length. We then generated a high-dimensional, acoustic parameterization of each 150 ms song window by taking the moving average in (across 20 ms windows segments every 5 ms) of seven song features (mean frequency, pitch, goodness of pitch, power 4, power 2 and summed log power of the spectrogram). We performed principal component analysis on this high-dimensional set across all bouts to normalize and reduce the feature set to 30 dimensions and applied the UMAP algorithm to project this high-dimensional representation into two dimensions^16,48^. The choice of parameters was made empirically to generate the clearest visual representation of the continuous bout trajectory^17,18^. This visualization gives a qualitative picture of the structure of the bout over a large set of renditions across the course of the degradation and recovery period. However, because the UMAP reduction algorithm makes nonlinear shifts in the distances between points across and within bouts, we do not use this visualization for quantitative analysis.

#### Duration distributions of best syllables

We tested whether the degradation and recovery duration pattern we found in our feedforward computational models after silencing a subset of neurons was present in our data. To do this, we considered the duration distributions of song segments over the course of the perturbation and recovery in our TenT experiment that most closely resembled the original stereotyped song (defined as the 2.5% of song segments on each day that were closest to the original stereotyped syllable, measured using the k-neighbor distance metric described above) (N = 4 birds). In this analysis we excluded stereotyped syllables which were either less than 100 ms in duration, never recovered, significantly altered their acoustic form, or did not segment cleanly into discrete clusters. In all four birds, we found syllable recovery trajectories that contained bimodal or multimodal duration distributions during days with degraded song (5/9 syllables analyzed).

### Song prevention

Adult zebra finches 120-150 dph (N = 3) were fitted with custom made fabric vests and a 20-40-gram weight attached to their vest, pulling them towards the ground to prevent them from acquiring singing posture. All birds received LV-TeNT injection bilaterally into HVC. Before we prevented the birds from singing, we allowed them to sing a few renditions to confirm that their songs were degraded. Afterward, the birds were restricted from singing for ∼10 days, during which they were monitored with a video camera to make sure they did not sing, although they could make calls. We confirmed that all the birds were able to move, perch, drink and eat freely and even allowed them to sing several renditions occasionally (1-3 every 2-3 days, which allowed us to track the degradation/recovery of songs). The size of the bullet weight on the birds had to be adjusted since they got accustomed to the weight in 2-3 days and attempted to sing more frequently. The weights and vest were removed daily, 1 hour prior to the light-off period while the experimenter stood next to the chamber to closely monitor that the birds were not singing. The vest and weights were put on the birds again as soon as the lights were turned back on the next day. After the prevention period, the birds were free to sing in their respective isolation chambers.

### Electrophysiology

Birds were first overdosed by an intramuscular injection of ketamine/xylazine (120/12 mg/kg) and after they became unresponsive to toe pinching, they were decapitated. The forebrain was quickly removed and kept in ice-cold slicing solution (in mM: sucrose 213, KCl 2.5, NaH_2_PO_4_ 1.2, NaHCO_3_ 25, Glucose 10, MgSO_4_ 2, CaCl_2_ 2, pH 7.4). Sagittal slices (300 μm) were cut using a vibratome (Leica VT1200S) and then incubated in HEPES holding solution (in mM: NaCl 102, KCl 2.5, NaH_2_PO_4_ 1.2 NaHCO_3_ 30, HEPES 20, Glucose 25, MgSO_4_ 2, CaCl_2_ 2, pH 7.35) at 34.5 °C for 30 minutes. Afterwards, slices were kept at room temperature (∼ 22 °C) between 30 minutes to 5 hours before being moved to the recording chamber. Bath ACSF (in mM: NaCl 124, KCl 2.5, NaH_2_PO_4_ 1.2, NaHCO_3_ 26, Glucose 25, MgSO_4_ 1, CaCl_2_ 2, pH 7.35, 33-34°C) was continuously perfused (∼2 mL/min) during recording. For current clamp whole-cell recordings, glass pipettes were filled with an intracellular solution (in mM): K-gluconate 135, MgCl_2_ 3, HEPES 10, EGTA 0.2, Na_2_-ATP 2, phosphocreatine 14, pH 7.25. HVC(RA) cells were identified by the presence of the retrograde fluorescent tracers injected into RA. Dual whole-cell recordings of HVC(RA) and HVC(INT) neurons were made between cells that were less than 100 µm apart. HVC(INT) neurons were identified by their relatively big soma, not having the retrograde fluorescent tracer, and spontaneous action potential firing^49^. We used voltage clamp to record miniature EPSCs with the following chemicals added to the bath (in µM): TTX 0.5, nimodipine 5 and picrotoxin 50, and glass pipettes were filled with (in mM) Cs(CH_3_)SO_3_ 135, MgCl_2_ 2, HEPES 10, EGTA 0.2, QX-314.Cl 5, Na_2_-ATP 2, phosphocreatine 14, pH 7.25. To record mIPSCs, the following chemicals were added to the bath (in µM): TTX 0.5, nimodipine 5, CNQX 10 and APV 25, and pipettes were filled with (in mM) CsCl 120, K-Gluconate 12, MgCl_2_ 2, HEPES 10, EGTA 0.2, QX-314.C1 5, Na_2_-ATP 2, phosphocreatine 14. For whole-cell NaChBac current recording, the bath solution did not contain any CaCl_2_ to eliminate Calcium currents, and TTX and 4-AP were added to block currents from the endogenous Na+/K+ channels, and glass pipettes were filled with (in mM) CsCl 135, MgCl_2_ 3, HEPES 10, EGTA 0.2, TEA-Cl 2, Na_2_-ATP 2, phosphocreatine 14. We only analyzed recordings in which access resistance was always smaller than 10 percent of the membrane resistance of the cell and no compensation was applied. Membrane potential was held at -70 mV to measure mEPSCs and -60 mV for mIPSCs. Liquid junction potential was not corrected. Electric signals were amplified and sampled at 20 kHz by an EPC-10 system (HEKA). Data analysis was performed off-line using Fitmaster (HEKA), Mini Analysis (Synaptosoft) and Matlab (Mathworks). Data were presented as mean ± s.e.m. Statistical difference was tested using one-way ANOVA followed by two-sided student’s t-test.

### Histology

After the experiments were concluded, animals were sacrificed, and their brains were processed for histological analysis. Animals were first deep anesthetized by intramuscular injection of ketamine/xylazine (100/10 mg/kg) and perfused intracardially with room temperature 3.2% PFA in 1xPBS. Brains were then extracted and incubated in the same fixative for 2-4 hours at room temperature. Each brain hemisphere was cut sagittally with a vibratome into 70-100 μm thick sections. The brain slices containing HVC were collected and incubated overnight with a rabbit anti-GFP antibody (EMD Millipore, AB3080P) in 1xPBS containing 10% donkey serum and 0.2% Triton at 4 °C. Sections were washed in 1xPBS with 0.05% Triton and incubated for 2 hours at room temperature with a secondary antibody (Abcam, ab150077). Brain slices were washed and mounted in Fluoromount (Sigma). Confocal images were taken with a LSM800 microscope. To validate the cell-type specificity of our LVs, we performed counterstaining in a subset of the brain slices, using known markers of inhibitory neurons, specifically anti-parvalbumin (Abcam, ab11427), anti-calretinin (SWANT, 7697) and anti-calbindin (SWANT, CB-300) (Extended Data Fig. 1b). Out of 1000 counted neurons, only one GFP+ cell was double-labeled with inhibitory markers.

### Modeling

We used network modeling to explore the role of different plasticity mechanisms.

#### Leaky Integrate and Fire Neurons

The membrane potentials *V*_*j*_ of neurons in all networks were modeled using leaky adapting integrate-and-fire dynamics:

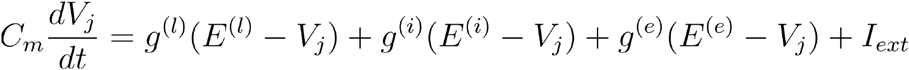

where *C*_*m*_=1e-6 F/cm^2^ is the membrane capacitance, *g*^(*l*)^,*E*^(*l*)^,*g*^(*e*)^,*E*^(*e*)^,*g*^(*i*)^,*E*^(*i*)^, the leak, excitatory, and inhibitory conductances and reversal potentials, and *I*_*ext*_ *∼ 𝒩* (0, *σ=*0.1 nA) a white noise current input. For excitatory neurons, *g*^(*l*)^*=*2.5e−2 S/cm^2^ and *E*^(*l*)^*=*−67 mV; for inhibitory neurons, *g*^(*l*)^*=*4e−2 S/cm^2^ and *E*^(*l*)^*=*−57 mV. For all neurons, *E*^(*e*)^ = 0 and *E*^(*i*)^ *=* —90 mV. When *V*_*j*_ reaches threshold, *V*_*th*_ =−43 mV the neuron spikes, and the voltage is reset to *V*_*re*_ (−67 mV for excitatory,−57 mV for inhibitory) after a refractory period *t*_*r*_ = 1 ms. The excitatory and inhibitory conductances are driven by incoming spike trains, represented by delta functions at times *t*_*k*_, from upstream neurons {*k*} filtered with a time constant *τ* = 4 mS:

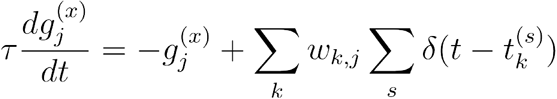

#### Network Architecture

We explored two different network architectures and the resulting dynamics. In the first network. we assumed all cells participate in the sequential dynamics. In the second network, we assumed that only a portion of the excitatory cells initially participate in the dynamics ^9^·30. To implement this, we assumed each neuron is silent with probability *p*_*s*_ = 0.4.

#### HVC Network

Following recent work, we modeled HVC as a feedfonvard, polychronous network^24,50^. The network is composed of 200 excitatory (E) and 50 inhibitory (I) neurons. In order to define the approximate difference in firing times between any neuron pair, we assign a coordinate *C*_*i*_ to each E cell, where *C*_*i*_ is the *i*^*th*^ sum of random variables uniformly distributed over [0,1]. Pairs of active excitatory cells are initially connected with a fixed weight *W* if the difference in their indices *C* is greater than zero and less than *c*\* = 40/3, ensuring feedforward propagation. Silent eel Is are assumed to be connected with in the network but with much lower probability, with no sensitivity to sequence order, and with cell-by-cell heterogeneity. Specifically, If one of a pair of neurons is silent, the cells are connected, with probability 0.075, with a weight *w*_*ij*_ drawn from an exponential distribution with a cell-specific mean *a*_*i*_ *W*, where *a*_*i*_ is uniformly distributed on [0,1]. For networks without silent neurons, *W* = 7e−5: for networks with 40% silent neurons, *W*= 4.9e−5. To implement recurrent inhibition observed in HVC ^51^, inhibitory neurons receive connections from excitatory cells with probability *P*_*ei*_ = 0.1 and weight *w*_*e,j*_ = 1e-4 and vice versa with probability *P*_*ie*_= 1 and weight *w*_*i,e*_ = 2*e−5*.

Following observations that axonal delays between HVC(RA) projectors are relatively long (1 - 7.5 ms)^24^and that HVC(RA) projectors typically synapse onto inhibitory interneurons close to their soma and other excitatory cells far from their soma ^52^, we implemented axonal delays in our model that reflected longer E-E axonal delays and relatively shorter E-1 and I-E delays. The delays between pairs of active E cells were *d*_*ij*_ = 3(*c*_*j*_ −*c*_*i*_) / ⟨*c*⟩] ms. When one of the pair was inactive, the delay was chosen randomly from a uniform distribution on [0, 3*c* * / ⟨*c*⟩] ms. This leads to E-E axonal delays that were 3 ms on average and roughly uniformly distributed. E ⟶ l and 1 ⟶ E delays were uniformly set to 0.5 ms. We found that a comparatively fast inhibitory pathway stabilized sequence dynamics by enabling inhibition to respond rapidly to changes in excitation.

#### Plasticity Rules

We then allowed networks to evolve under both firing rate homeostasis and spiking timing-dependent plasticity. All synapses subject to plasticity were given a lower bound *w*_*min*_. Af1er each trial, synaptic strength *w*_*i,j*_ evolved according to

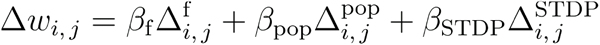

where weights are updated due to single-cell firing rate homeostasis (Δ^f^), local activity homeostasis (Δ ^pop^) and STDP (Δ ^STDP^) respectively, and *β*_f_*=* 0.06,. *β*_pop_ = 0.01, and. *β*_STDP_ = le-4 *β*_f_ was chosen such that firing rate homeostasis could bound potentiation due to STOP. *β*_pop_ was chosen to be small so that recruitment of formerly silent neurons would occur slowly. All E-E synaptic strengths had upper bound 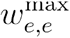,all 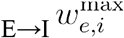.

##### Firing rate homeostasis

Single-cell firing rate homeostasis moves a neuron’s firing rate toward a set point a set point 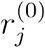 according to:

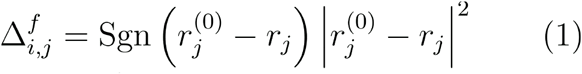

where ^*r*^_*j*_ is the average firing rate of neuron *j* in that trial (refer to Figure 4).

##### Local population activity homeostasis

Taking inspiration from literature that has shown homeostasis may operate on a network scale, we included in our final model a form of homeostasis that permits individual neurons to monitor and respond to the activity of their neighbors. We implemented here one potential mechanism by which such local population activity homeostasis might be achieved, based loosely on the TNFa pathway 38. We assume each E neuron secretes a chemical factor which diffuses locally in space, the concentration of which is proportional to the neuron’s own activity level. All E neurons are assumed to monitor the local concentration of this factor and adjust their incoming excitatory synapses in order to maintain a target concentration. To implement this form of local population homeostasis, each excitatory neuron was first assigned a location by uniformly sampling the space within a unit sphere. The local factor concentration that an excitatory neuron senses is then

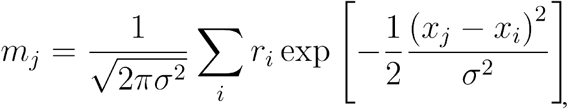

Where *x*_*j*_ is the location of neuron *J*, and *σ* is a parameter controlling the spatial extent of each neuron’s diffuse release (*σ*^2^ = 0.3). The corresponding update to local population homeostasis is

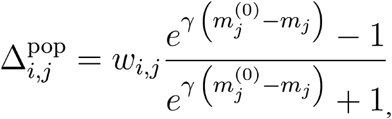

where 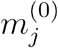 is the local concentration setpoint of neuron *j* and *γ* = 0.01 dictates the strength of the local population homeostasis near the setpoint. Local concentration setpoints were chosen by computing the average local concentration over 50 activations of the network prior to the perturbation.

In the version of our model that included local population homeostasis and inactive neurons, we modified single-cell firing rate homeostasis in E neurons to act only to decrease input if firing exceeded a neuron’s setpoint under the assumption that connectivity (and not homeostatic properties) should dictate whether a neuron was active or inactive. The modified rule is given by

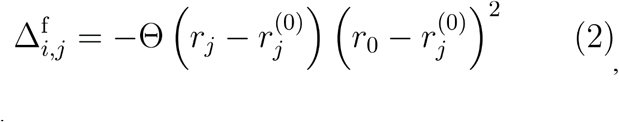

where Θ is the Heaviside function.

##### Hebbian plasticity

We found that an antisymmetric pairwise STDP rule with a reasonable time constant (20 ms) did not maintain the relative firing times of a sequence when the spike trains of successively firing neurons overlapped. A burst timing dependent plasticity (BTDP) rule ^50^ that low pass filtered spike trains before applying pairwise STDP was able to maintain relative firing times, but could lead to contraction of the sequence if noise enabled neurons to spike earlier than their typical firing times. In contrast, we found that a triplet STDP rule well maintained the firing times of neurons in a feedforward, excitatory network. The minimal triplet rule introduced by Pfister and Gerstner depends on triplets of spikes (post-pre-post) for potentiation and pairwise interactions for depression ^53^. The update for the triplet rule for excitatory neurons was

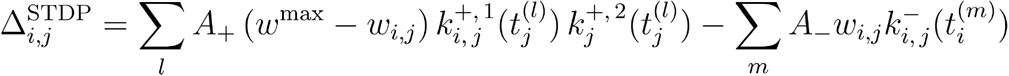

where *w*^max^ differs for E →E and E →I synapse, *m* and *l* index the spike times of the pre- and postsynaptic neuron, respectively, and the *k* variables keep track of relative spike timing and implement the eligibility window:

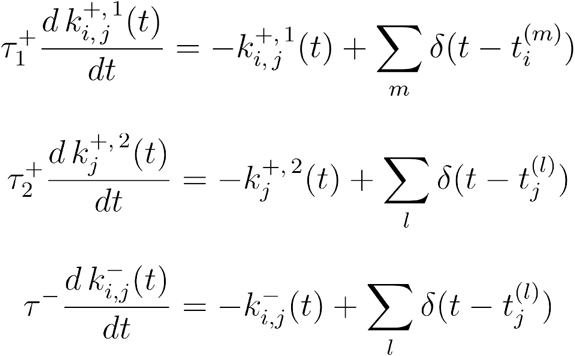

The constants regulating the relative strength of potentiation and depression, *A*_+_= 6.5 and *A*_−_ = 7.1(250 and 0, respectively, for E → I synapses), and the timescales of STDP, *τ*_−_ = 33.7, 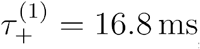 and 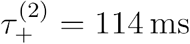, were inspired by the parameters given by Pfister and Gerstner in their triplet STDP model of neurons in the visual cortex. We found sequences were most stable when when potentiation due to triplets was implemented in “nearest spike” spike fashion, i.e. 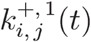 and 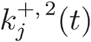 were bounded by [0, 1]. To stabilize E →I STDP, each inhibitory neuron was assigned a total excitatory synaptic input bound determined by the neuron’s total excitatory synaptic input at the beginning of the simulation. When the bound was exceeded, E-I weight *w*_*ij*_ was rescaled as

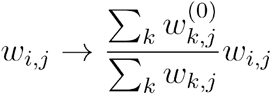

where 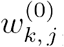 represents the size of the synapse prior to the first activation of the network. No plasticity takes place on I-E connections.

#### Activation of Networks

At the beginning of each trial, networks were active for 10 ms, after which each of the first 10 neurons of the network was driven by an independent burst (4 spikes, 660 Hz), with onset times drawn from a Gaussian (mean 10 ms, STD 1 ms). Input weights were chosen to produce reasonable spiking behavior in the first layers of the network. Networks were simulated for an additional 90 ms following the stimulus. The time step for all simulations was 0.1 ms.

#### Network Initialization Procedure

To initialize networks, an activation of each network was first simulated once without plasticity. Neurons that fired during this trial were assumed to be active. Active neurons were assigned a uniform firing rate setpoint (*r*^(0)^ *=* 4) and were subject to cell-autonomous firing rate homoeostasis as given in Eq. (1) and STDP for 500 additional activations of the network. For networks with population homeostasis, {*m*_*i*_}, the local secreted factor concentrations were computed for activations 500-550, and then averaged to set 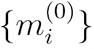 from this point, population homeostasis was permitted to act on the network, and the cell-autonomous firing rate homeostasis rule was changed to Eq. (2).

#### Simulated Tetanus Toxin Perturbation of Neurons

Tetanus toxin perturbation of the network was simulated by randomly selecting a cell with probability *P*_*T*_ for perturbation and removing all its outgoing connections. Perturbed cells were assumed to not contribute to the population activity level.

#### Simulated Tetanus Toxin Perturbation of Neurons

Tetanus toxin perturbation of the network was simulated by randomly selecting a cell with probability for perturbation and removing all its outgoing connections. Perturbed cells were assumed to not contribute to the population activity level.

Networks were then allowed to evolve for 4000 renditions according to the plasticity rules described above. We classified networks as ‘recovered’ if for >90% of renditions 4000 to 4100 (i) any neuron fired in each of the following ranges of cell indices: 0-20, 100-120, and 180-200, (ii) the average spike time of cells 0-20 preceded that of cells 100-120, etc., and (iii) network activity persisted for >65 ms after the onset of activation (typical network sequence duration was 80 ms initially). be presented together with its figure. Also, please include line numbers within the text.

