## Supplemental Data for "Unsupervised Restoration of a Complex Learned Behavior After Large-Scale Neuronal Perturbation"

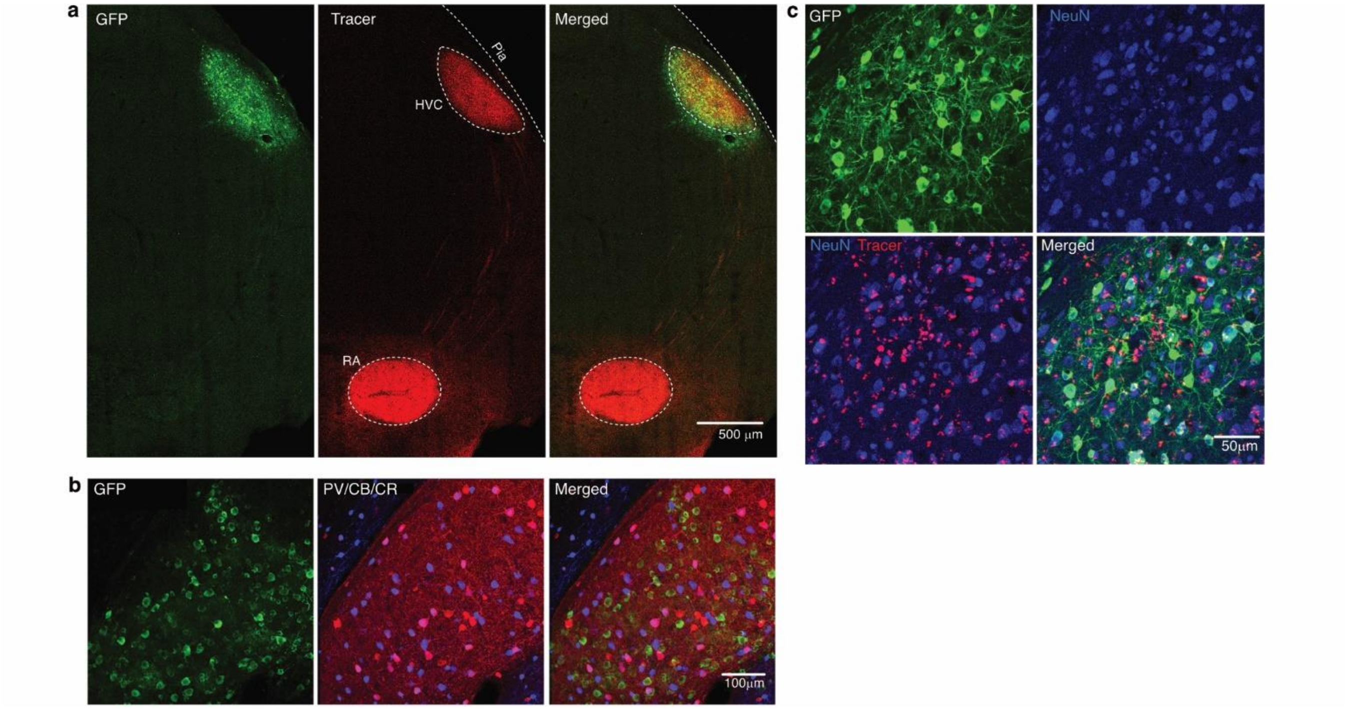


**Extended Data Figure 1 Specific infection of HVC projection neurons by LVs**

**(a)** Confocal image of a brain slice from a bird injected with LVs, showing the expression of the transgene (tagged with GFP) in HVC, which is labeled by a fluorescent retrograde tracer injected into RA; **(b)** Confocal images of a brain slice showing that LV selectively target projection neurons, where the LV transgene (tagged with GFP, seen in green) does not overlap the immunofluorescent signal of pooled antibodies against some of the standard markers of inhibitory neurons (PV, parvalbumin/CB, calbindin/CR, calretinin, seen in red, blue); **(c)** Confocal images showing the expression of delivered transgene (tagged with GFP) in the majority of HVC RA-projecting neurons (retrogradely labeled with fluorescent RetroBeads from RA).

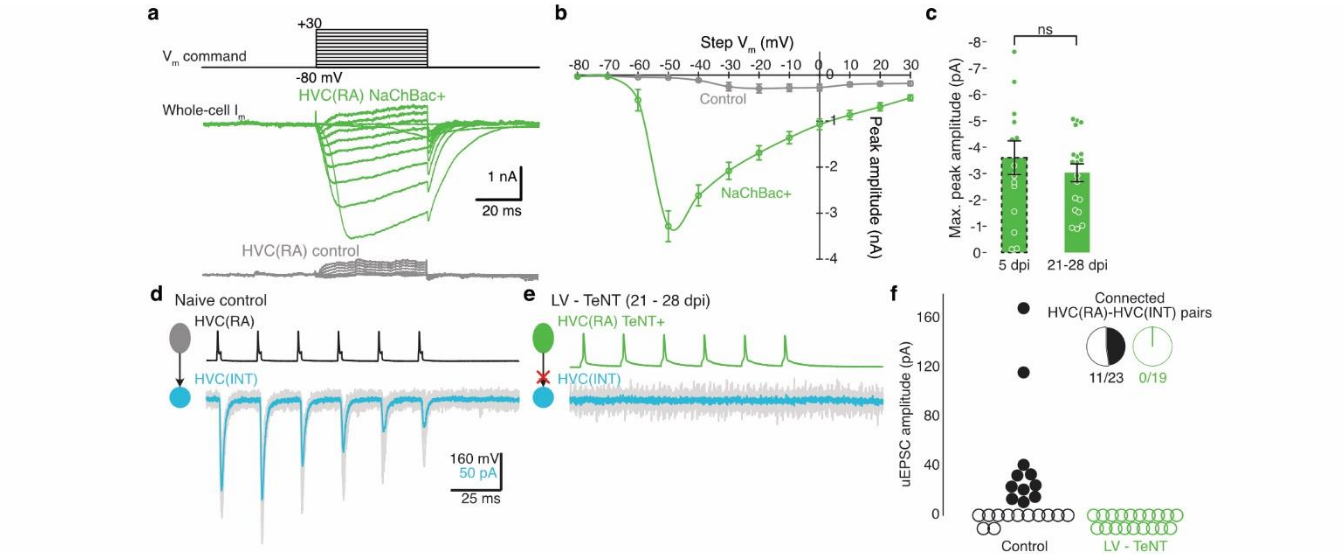


**Extended Data Figure 2 Stable expression of transgenes in HVC projection neurons**

**(a)** Example traces of whole-cell NaChBac current evoked by depolarizing voltage steps (from -80 to +30 mV. increment, 10 mV.) recorded from HVC(RA) neurons infected with LV-NaChBac (NaChBac+) or control cells; **(b)** I-V curve of the NaChBac current, illustrating the peak amplitude of whole-cell current at different step voltages; **(c)** Comparison of the maximal peak amplitude of whole-cell NaChBac currents recorded at 5 dpi (3.6 ± 0.6 nA) vs. 21-28 dpi (3.0 ± 0.3 nA), demonstrating that the presence of NaChBac is stable, as it could be detected even >3 weeks after viral injection, after the song had already recovered. Student's t-test. N = 10/3 (control), 15/2 (5 dpi), and 19/3 (21-28 dpi). Error bars represent s.e.m.; **(d)** Example traces from a dual patch clamp recording, showing that when action potentials were evoked in RA-projecting HVC (HVC(RA)) neurons (upper), excitatory postsynaptic currents (EPSCs) can be reliably detected in the connected interneuron (HVC(INT)) (below); **(e)** Example traces from dual patch clamp recording made between an HVC(RA) neuron infected with LV-TeNT and an interneuron, showing that no EPSC could be detected in the interneuron when the HVC(RA) neuron was stimulated; **(f)** Summary of dual patch clamp results from LV-TeNT neurons. In control animals, 11 out of 23 recorded pairs were connected (the amplitude of EPSCs shown in the scatter plot), but none of the 19 pairs recorded between TeNT+ projection neurons (21-28 dpi) and neighboring interneurons was connected. The block of synaptic transmission by TeNT expression is long-lasting, as it could be detected even >3 weeks after viral injection, after the song had already recovered.

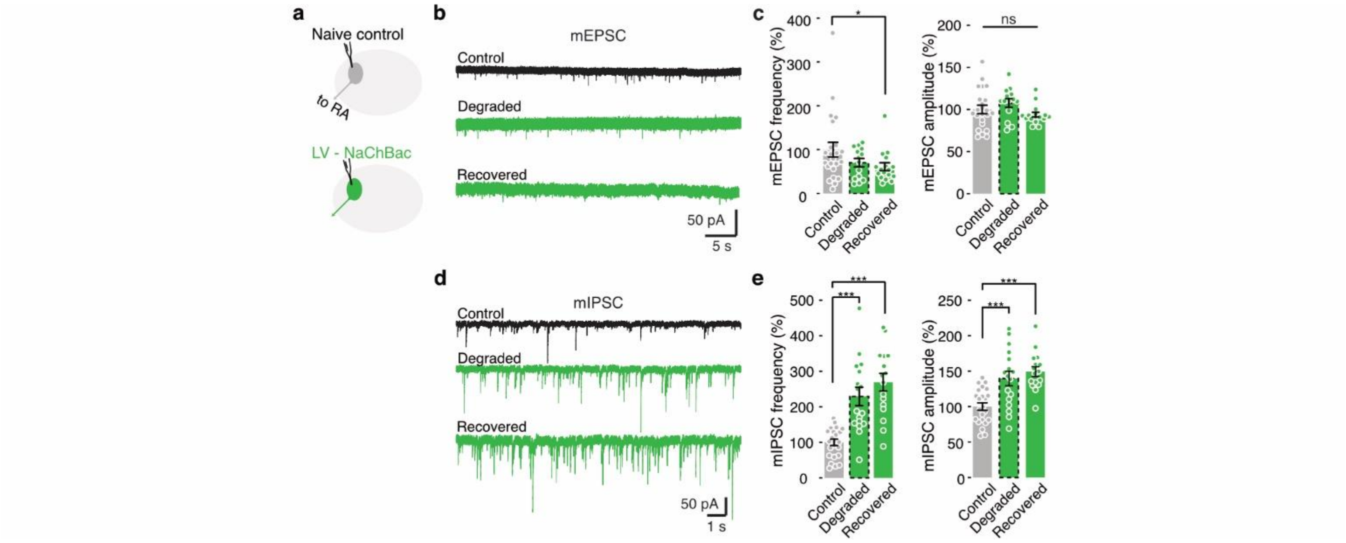


**Extended Data Figure 3 Changes in synaptic currents in NaChBac+ HVC(RA) cells**

**(a)** Schematic drawing showing whole-cell patch clamp recordings were made in HVC(RA) neurons infected with LV-NaChBac or naive controls; **(b)** Example of current traces with mEPSC events recorded in HVC(RA) neurons expressing NaChBac at different time points. "Degraded" illustrates currents at 5 dpi, when the song was highly irregular. "Recovered" illustrates currents at 25-35 dpi, after the song was fully recovered; **(c)** Group data of the frequency and amplitude of mEPSCs in HVC(RA) NaChBac+ cells. mEPSC frequency: Control, 9.7 ± 1.6 $min^{-1}$, N = 22/4; Degraded, 6.8 ± 0.9 $min^{-1}$, N = 14/5; Recovered, 5.9 ± 0.9 $min^{-1}$, N = 17/4. mEPSC amplitude: Control, 17.2 ± 0.9 pA; Degraded, 18.5 ± 0.9 pA; Recovered, 16.1 ± 0.5 pA; **(d)** Example of current traces with mIPSC events recorded in HVC(RA) neurons expressing NaChBac at different times after injection; **(e)** Group data of the frequency and amplitude of mEPSCs in HVC(RA) NaChBac+ cells. mIPSC frequency: Control, 2.7 ± 0.2 $s^{-1}$, N = 23/4; Degraded, 6.1 ± 0.7 $s^{-1}$, N = 17/3; Recovered, 7.2 ± 0.7 $s^{-1}$, N = 16/5. *, $p < 0.05$, ***, $p < 0.001$, student's t-test. Error bars represent s.e.m..

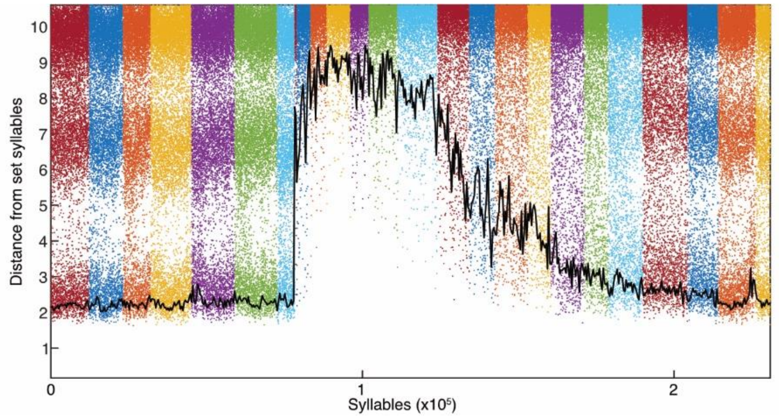


**Extended Data Figure 4 Tracking the degradation and recovery of song syllables using k-neighbor acoustic distance from set syllables**

(See also the Methods for a detailed explanation). Here we plot the distance of each analyzed syllable to the set of the syllables in the original song, in a high dimensional space constructed using multiple acoustic features. Each dot represents one syllable, and they are chronologically ordered on the x axis. Different colors are used to denote different days. The black line represents the average k-nearest neighbor distance of the closest song segments (2.5%) to the original syllable set, which illustrates the dynamic trajectory of song degradation and recovery. We tracked the change of songs in the same way in Figs. 2e, 4d, and Extended Data Figs. 5c, 8c.

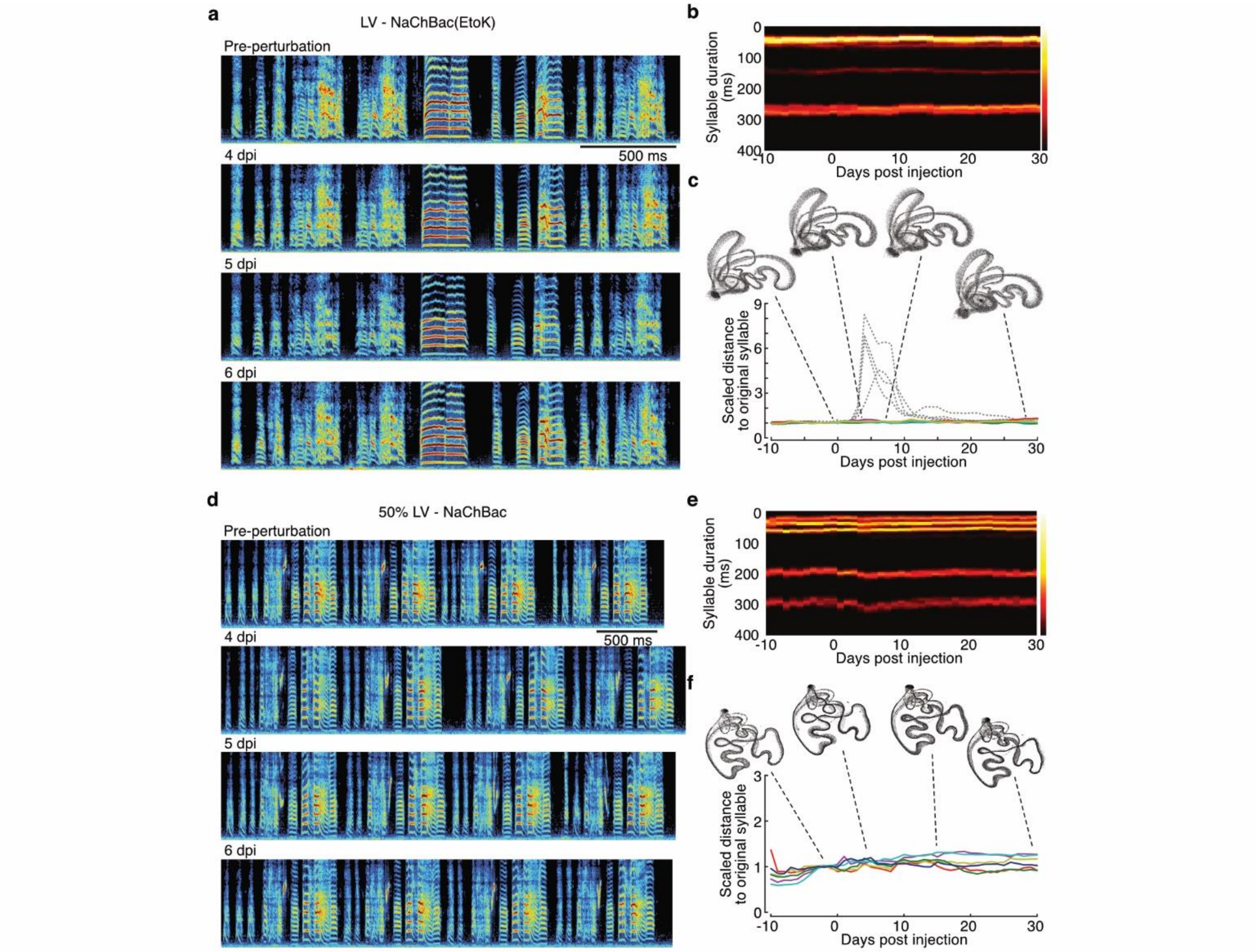


**Extended Data Figure 5 Songs degradation and recovery was not due to mechanical lesion or inflammation**

**(a)** Example spectrograms of songs from a bird injected with LV-NaChBac(EtoK), a ded-pore mutant of NaChBac; **(b)** Distribution of syllable durations per day of the same bird shown in panel a; **(c)** Plots of scaled acoustic distance to original syllables. Data was from the same bird as in panels a and b. The insets are UMAP visualizations of songs at selected time points. Dashed lines are generated from the syllables of the bird injected with LV-NaChBac shown in Fig. 1d, and for comparison here they were not normalized to maximum. Experiments with LV-NaChBac(EtoK) yielded consist results, N = 4; **(d)** Example spectrograms of songs from a bird injected with half of the volume of LV-NaChBac used for animals shown in Fig. 1; **(e)** Distribution of syllable durations per day of the same bird shown in panel d; **(f)** Plots of scaled acoustic distance to original syllables. Data was from the same bird as in panels d and e. Each line/color represents one syllable. The insets are UMAP visualizations of songs at selected time points. Experiments with reduced volume of LV-NaChBac were repeated, N = 2.

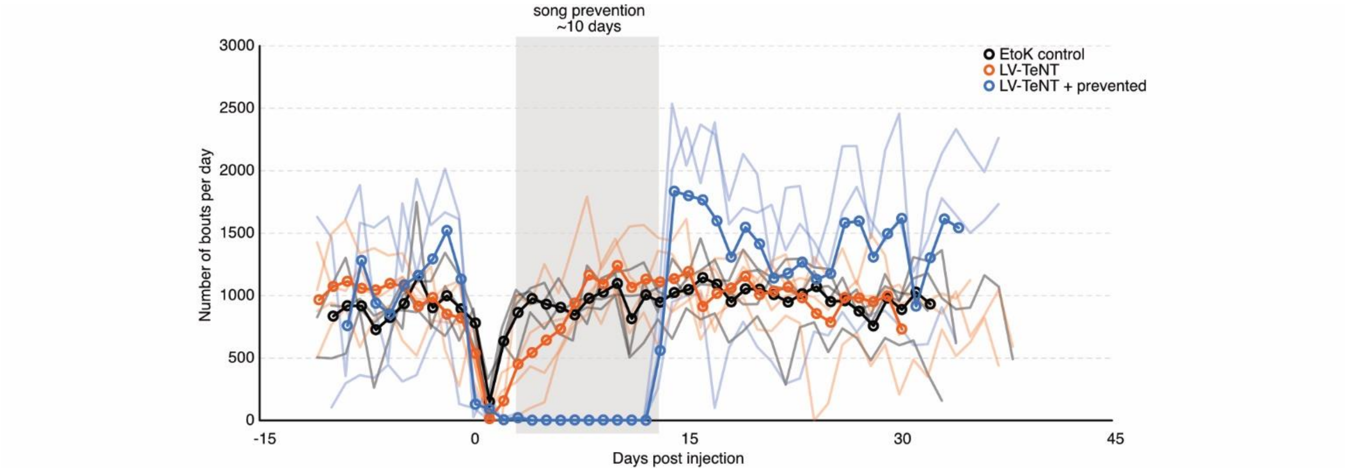


**Extended Data Figure 6 Number of song bouts sung per day in freely signing animals and individuals prevented from singing**

A song bout was defined as a continuous vocalization without a gap longer than 450 ms, and it usually includes several motifs. Faded lines are numbers of song bouts per day of each individual bird. Bold lines are the mean value per day of each group. Song prevention period is marked by a gray square. Notice that in cases, animals sing very few bouts on the day after the surgical procedure.

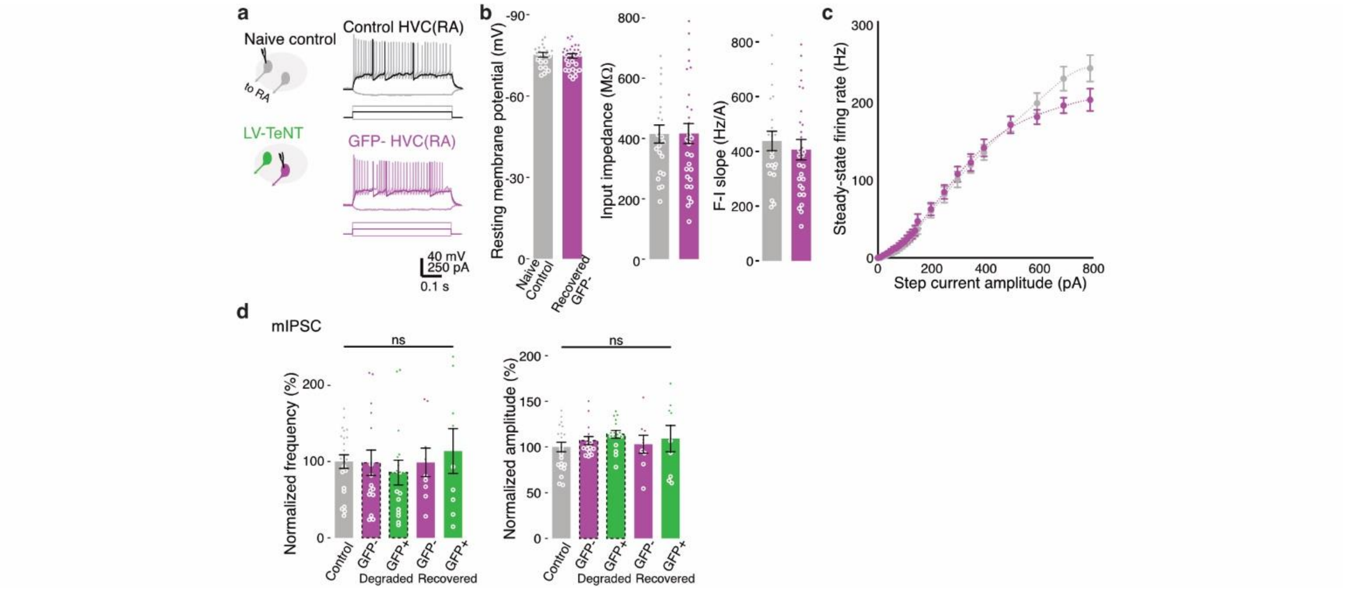


**Extended Data Figure 7 Unperturbed cells did not change their intrinsic excitability or inhibitory synaptic inputs**
**(a)** (Left) Whole-cell patch clamp recordings were made in GFP- HVC(RA) neurons in naive control birds or birds injected with LV-TeNT. (Right) Membrane potential and firing pattern of HVC(RA) neurons in response to current steps; **(b)** Group data showing that no significant difference was found in the resting membrane potential (-75.2 ± 0.8 vs. -74.8 ± 0.8 mV), input resistance (414.8 ± 30.0 vs. 416.2 ± 33.1 MOhm), or initial F-I slope (437.9 ± 35.5 vs. 406.2 ± 37.3 Hz/A) between neurons in naive control (N = 22/3) and in birds with LV-TeNT for more than 25 days (N = 30/4). Student's t-test; **(c)** F-I curves obtained from HVC(RA) neurons showed no difference between control or animals recovered from LV-TeNT; **(d)** Group data of mIPSC recorded in HVC(RA) neurons, showing no significant difference at any time during the experiment. "Degraded" indicates the time when the song was degraded, at 5 dpi. "Recovered" indicates the time after the song had fully recovered, at 25 dpi. mIPSC frequency: Control, 2.7 ± 0.2 $s^{-1}$, N = 23/4; "Degraded" GFP-negative cell, 2.6 ± 0.4 $s^{-1}$, N = 16/3; "Degraded" GFP-positive, 2.3 ± 0.4 $s^{-1}$, N = 16/3; "Recovered" GFP-, 2.6 ± 0.5 $s^{-1}$, N = 9/3; "Recovered" GFP+, 3.1 ± 0.8 $s^{-1}$, N = 9/3. mIPSC amplitude: Control, 37.9 ± 2.0 pA; "Degraded" GFP-negative cell, 40.6 ± 1.7 pA; "Degraded" GFP-positive cell, 43.2 ± 1.6 pA; "Recovered" GFP- negative cell, 39.0 ± 3.7 pA; "Recovered" GFP- positive cell, 41.4 ± 5.4 pA. One-way ANOVA. Error bars represent s.e.m.

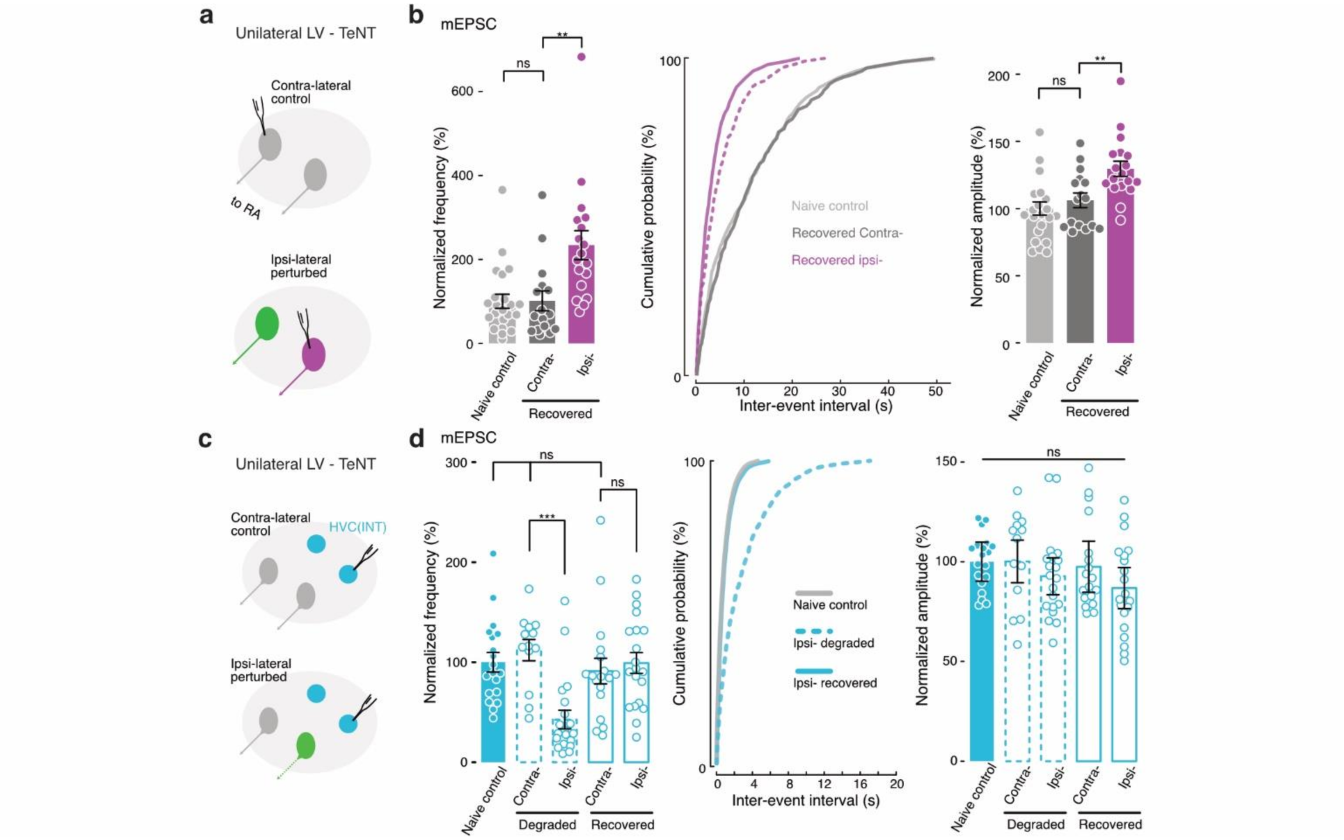


**Extended Data Figure 8 Synaptic changes in unmanipulated neurons only occurred in the injected hemisphere**
**(a)** Whole-cell recordings were made in GFP- HVC(RA) ("unmanipulated") neurons in the injected and unperturbed hemispheres of birds with unilateral LV-TeNT injection; **(b)** Group data of mEPSC recorded in naive control birds and birds with unilateral LV-TeNT. The frequency (left) and amplitude (right) of mEPSC increased significantly in GFP- HVC(RA) neurons in the injected side, as compared with neurons in the contralateral side or naive animals. (Middle) Cumulative curve of inter-event intervals of mEPSCs. Dashed line represents data from GFP- neurons from animals with bilateral LV-TeNT injection for comparison and was adapted from Figure 4c. mEPSC frequency: Naive control, 9.7 ± 1.6 $min^{-1}$, N = 23/4; Contralateral (uninjected HVC), 9.8 ± 2.3 $min^{-1}$, N = 16/4; Ipsilateral (injected), 22.8 ± 3.3 $min^{-1}$, N = 18/3. mEPSC amplitude: Naive control, 17.2 ±0.9 pA; Contralateral, 18.2 ± 0.9 pA; Ipsilateral, 22.3 ± 1.0 pA; **(c)** Whole-cell recordings were made in inhibitory (HVC(INT)) neurons in birds with unilateral LV-TeNT; **(d)** Group data of mEPSC recorded in birds with LV-TeNT at 5 dpi ("Degraded") or > 25 dpi ("Recovered"). (Left) The frequency of mEPSC in HVC(INT) neurons decreased after virus injection, but eventually recovered to a level comparable to that of controls. (Middle) Cumulative curve of inter-event intervals of mEPSCs. mEPSC frequency: Naive control, 1.47 ± 0.14 $s^{-1}$, N = 19/2; Degraded contralateral, 1.56 ± 0.16 $s^{-1}$, N = 13/3; Degraded ipsilateral, 0.63 ± 0.14 $s^{-1}$; N = 20/3; Recovered contralateral, 1.34 ± 0.19 $s^{-1}$, N = 18/4; Recovered ipsilateral, 1.45 ± 0.15 $s^{-1}$, N = 20/4. mEPSC amplitude: Naive control, 38.3 ± 1.2 pA; Degraded contralateral, 38.4 ± 2.6 pA; Degraded ipsilateral, 35.5 ± 1.9 pA; Recovered contralateral, 37.3 ± 2.1 pA; Recovered ipsilateral, 33.2 ± 2.0 pA. **, $p < 0.01$, ***, $p < 0.001$, ANOVA & student's t-test. Error bars represent s.e.m.

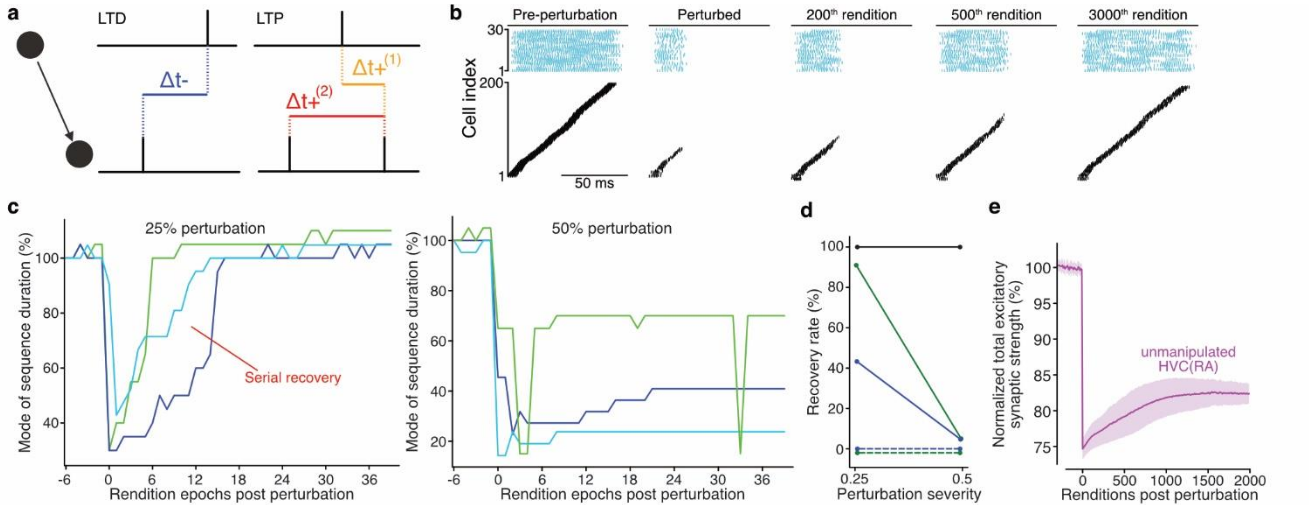


**Extended Data Figure 9 Recovery of modeled sequences with STDP**

**(a)** Schematic showing triplet STDP implemented both in E→E and E→I synapses. Potentiation is mediated by triplets of spikes; depression is dictated by a pairwise rule. See Methods for details; **(b)** Spike raster plots showing the sequential dynamics generated by HVC neurons before and after perturbation with only STDP and downward firing rate homeostasis (see Eq. 2 in Methods) implemented; **(c)** Plots of the normalized mode of sequence duration (computed for epochs of 50 renditions) against epoch number when 25% (left) or 50% (right) of HVC(RA) neurons are perturbed in networks with E→E STDP and downward firing rate homeostasis. Note that when 25% of neurons were lost, most networks exhibited serial regrowth of their sequence durations and eventually recovered. When 50% of neurons were perturbed, E→E STDP was largely unable to restore sequential dynamics; **(d)** Fraction of networks that recovered sequential activity under various plasticity rules after 25% and 50% perturbations. Networks with strong E→E STDP (green) recovered at a high rate (24 of 25 networks recovered) when 25% of neurons were perturbed, but generally did not recover when 50% of neurons were perturbed (1 of 25 networks recovered) or when E→I STDP was introduced. Dashed lines represent networks with E→I STDP (see Methods, Hebbian plasticity. 0 of 25 networks recovered). When E→E STDP was weakened (blue lines), the recovery rate fell (11 of 25 networks recovered). When the coefficient of E→E STDP was increased above the value for networks shown in green, networks were generally unstable. Recovery rates of networks with STDP and full firing rate homeostasis shown in black; **(e)** Plot of the normalized total excitatory synaptic input per unmanipulated HVA(RA) neuron against renditions post 25% perturbation for networks with E→E STDP and downward firing rate homeostasis. Average excitatory synaptic weight for recovered networks never exceeded the pre perturbation value.

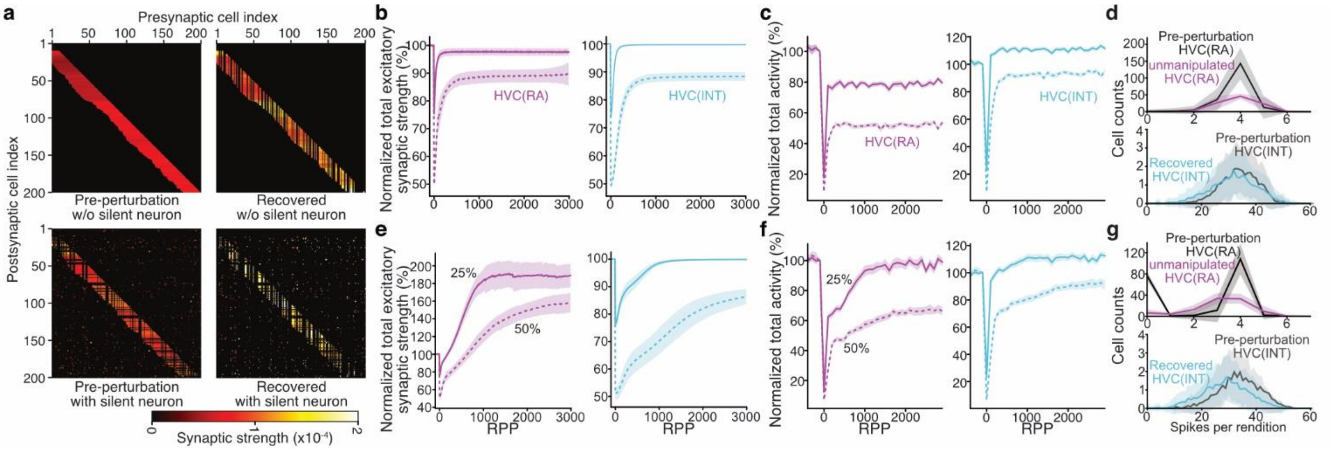


**Extended Data Figure 10 Recovery of modeled sequences with population-level homeostatic plasticity**

**(a)** E→E synaptic connectivity matrices in networks without (top) and with (bottom) silent neurons both pre perturbation (left) and following recovery (right). STDP, downward firing rate homeostasis, and population homeostasis rules were implemented. Note in the bottom right panel that the population rule strengthens the diffuse, off-diagonal connectivity to silent neurons, increasing the probability they will fire; **(b-d)** are generated using networks with firing rate homeostasis, STDP, and local population homeostasis but without silent neurons; **(e-g)** are generated when silent neurons are added; **(b,e)** Normalized average total excitatory input into each unmanipulated HVC(RA) neuron (purple) and interneurons (blue) for 25% (solid lines) and 50% (dashed) perturbations. Note with silent neurons, an overshoot in E→E weight was introduced for both 25% and 50% perturbations; **(c,f)** Normalized total activity of all functional HVC(RA) (purple) and HVC(INT) (blue) neurons per rendition. Note the addition of silent neurons is able to improve the fraction of initial excitatory activity recovered; **(d,g)** Distributions of firing rate per rendition for HVC(RA) (top) and HVC(INT) (bottom) neurons following 50% perturbation. Pre-perturbation distributions in black or gray. RPP, rendition(s) post perturbation. Shadows in all panels represent standard error.
